# Different representations in layer 2/3 and layer 5 excitatory neurons of the primary visual cortex

**DOI:** 10.1101/2025.06.30.662366

**Authors:** Timothy P.H. Sit, Célian Bimbard, Anna Lebedeva, Matteo Carandini, Philip Coen, Kenneth D Harris

## Abstract

The cortex contains multiple types of excitatory neuron, differentiated primarily by their layer of residence. We recorded from neuronal populations in the mouse visual cortex using 2-photon calcium imaging or Neuropixels probes, and found that excitatory neurons in layer 2/3 (L2/3) and layer 5 (L5) differed in their encoding of visual vs. nonvisual signals, with L2/3 more strongly modulated by visual stimuli and L5 more strongly modulated by movement. Movement had opposite effects on population synchrony in the two layers: it desynchronizes L2/3, by abolishing spontaneous synchronous fluctuations that entrain that population, and synchronizes L5, where excitatory cells are less entrained by spontaneous synchronous fluctuations and more strongly correlated with movement itself. Spontaneous activity was lower-dimensional in L2/3 than L5, with L2/3 population activity dominated by a single dimension of overlap between spontaneous and stimulus-evoked subspaces. We conclude that excitatory neurons in different layers of the visual cortex carry different representations of visual and non-visual signals.

## Introduction

Neuronal populations in sensory cortex exhibit complex patterns of spontaneous activity, as well as sensory responses^1^. The structure of cortical spontaneous activity shares at least some features with sensory responses: widefield imaging has shown similarities in the spatiotemporal structure of spontaneous and evoked activity patterns across the cortical surface^2–5^; and electrode recordings in multiple species and sensory areas have found similarities in spontaneous and evoked correlation patterns^6–10^. This similarity has led to computational interpretation: that spontaneous firing patterns comprise recollections of previous events or samples from a learned generative model of its sensory inputs^11–16^.

More recently, large-scale recordings in awake animals have suggested a second perspective: spontaneous cortical activity at least in part reflects ongoing behavior. Activity in sensory cortex is modulated by behaviors such as running or whisking even in the absence of direct sensory input and spontaneous movements and related state variables explain a large fraction of the variance of population activity across brain areas^17–24^. This modulation has been suggested as a form of integration into a high-dimensional mixed representation of sensory and motor variables^22,23,25,26^, or as predictions of sensory input due to self-motion, which can then be subtracted from actual sensory input, in a manner that may differ between layers^24,27–31^.

Furthermore, two-photon imaging has suggested that, at least in L2/3 of mouse primary visual cortex, spontaneous activity and sensory responses overlap almost entirely along a single dimension^22^. If the spontaneous–evoked overlap were truly one-dimensional, this would challenge the interpretation of spontaneous activity as an internal model of sensory stimuli: although one dimension of overlap is sufficient to generate statistically significant similarity of spontaneous and evoked population patterns^32^, one dimension of overlap is not sufficient to capture differences between recalled stimuli or to encode a rich representation of the sensory world.

A possible resolution of these apparently contradictory results would be if the relationship between spontaneous and evoked activity depends on the cortical population sampled, and on cortical state. Extracellular electrophysiology is biased toward higher firing-rate neurons found in deep cortical layers, which are easier to spike sort than more numerous but sparsely active superficial neurons^33^. 2-photon calcium imaging has an opposite bias to sparsely-active neurons in superficial layers. Furthermore, cortical population activity has a fundamentally different character in different behavioral states: in alert behaviors, cortical spontaneous activity is relatively steady (the “desynchronized state”), but during behavioral quiescence or sleep the cortex enters a “synchronized state”, where large coordinated fluctuations invest population activity^34^. The effects of state seem to differ between layers^35^, and layer-specific motion signals have been reported in visual, auditory, and somatosensory cortices^36–39^.

Here we revisit the similarity of spontaneous and evoked activity by directly comparing excitatory populations in superficial and deep layers of mouse V1, using both two-photon calcium imaging and laminar Neuropixels recordings, during stationary and movement states. We find fundamental differences in the structure of spontaneous and evoked activity between layers: spontaneous activity in deep layers more strongly encode movement signals; superficial layers more strongly encode sensory responses but their spontaneous activity becomes dominated by 1-dimensional synchronized-state fluctuations during behavioral quiescence. In all behavioral states, the overlap of spontaneous and evoked activity was almost exclusively 1-dimensional in L2/3, but less so in deep layers.

## Results

To investigate the differences in the encoding of sensory-evoked and spontaneous activity between cortical layers, we performed two-photon calcium imaging of visual cortical populations in two mouse lines. To record from layer 2/3 (L2/3) excitatory neurons we used the TetO-GCaMP6s;CaMKIIa-tTA mouse line^40^, which labels all excitatory cells, resulting in good recordings of superficial layers but occlusion of deep layers. To record from deeper layers, we crossed Rbp4-Cre KL100^41,42^ with Ai94 (TITL-GCaMP6s)^43^, which labels layer 5 (L5) excitatory neurons without occlusion because more superficial cells are not labelled. In each session, we recorded spontaneous activity (gray-screen conditions) and responses to combinations of natural videos and sounds, as previously described^44^. We monitored the orofacial movement of mice videographically during the experiments. In addition to recording large populations with 2-photon microscopy, we analyzed the responses of smaller populations recorded with Neuropixels probes in response to the same stimuli.

### Movement correlates are more uniform in L5 than L2/3

The correlation of V1 activity with movement was stronger and more positive in L5 than L2/3. We computed each neuron’s Pearson’s correlation with videographic motion energy during the spontaneous period, when only a gray background is presented on the three screens. Consistent with previous results^22^, in L2/3 we observed both positive and negative correlations with movement energy. By contrast, in L5 correlations were larger and more commonly positive (Figure 1A, B). The negative correlations with movement in L2/3 were genuine: a linear shift test^45^ revealed that of L2/3 cells that significantly correlated with movement, 61% ± 9% (mean ± s.e.m, N = 18 sessions) correlated positively and the remainder negatively. However in L5 almost all significant correlations were positive (97% ± 2%; mean ± s.e.m, N = 16 sessions; Figure S1). Correlations (re-gardless of significance) were stronger in L5 (i.e. larger absolute value; Fig. 1C; p = 0.02, Rank sum test, N = 18 and 16 sessions in L2/3 and L5) and a larger fraction of correlations (whether significant or not) were positive in L5 than L2/3 (Figure 1D; p = 0.003, Rank sum test, N = 18 and 16 sessions in L2/3 and L5). Analysis of Neuropixels recordings in V1^44^ showed similar results (Figure S3A-D), and showed that this effect of larger and more positive correlation in L5 cells persists even accounting for differences in firing rate between L2/3 and L5 cells (Figure S8A, C, D) and across multiple timescales (Figure S8E, F).

**Figure 1.**
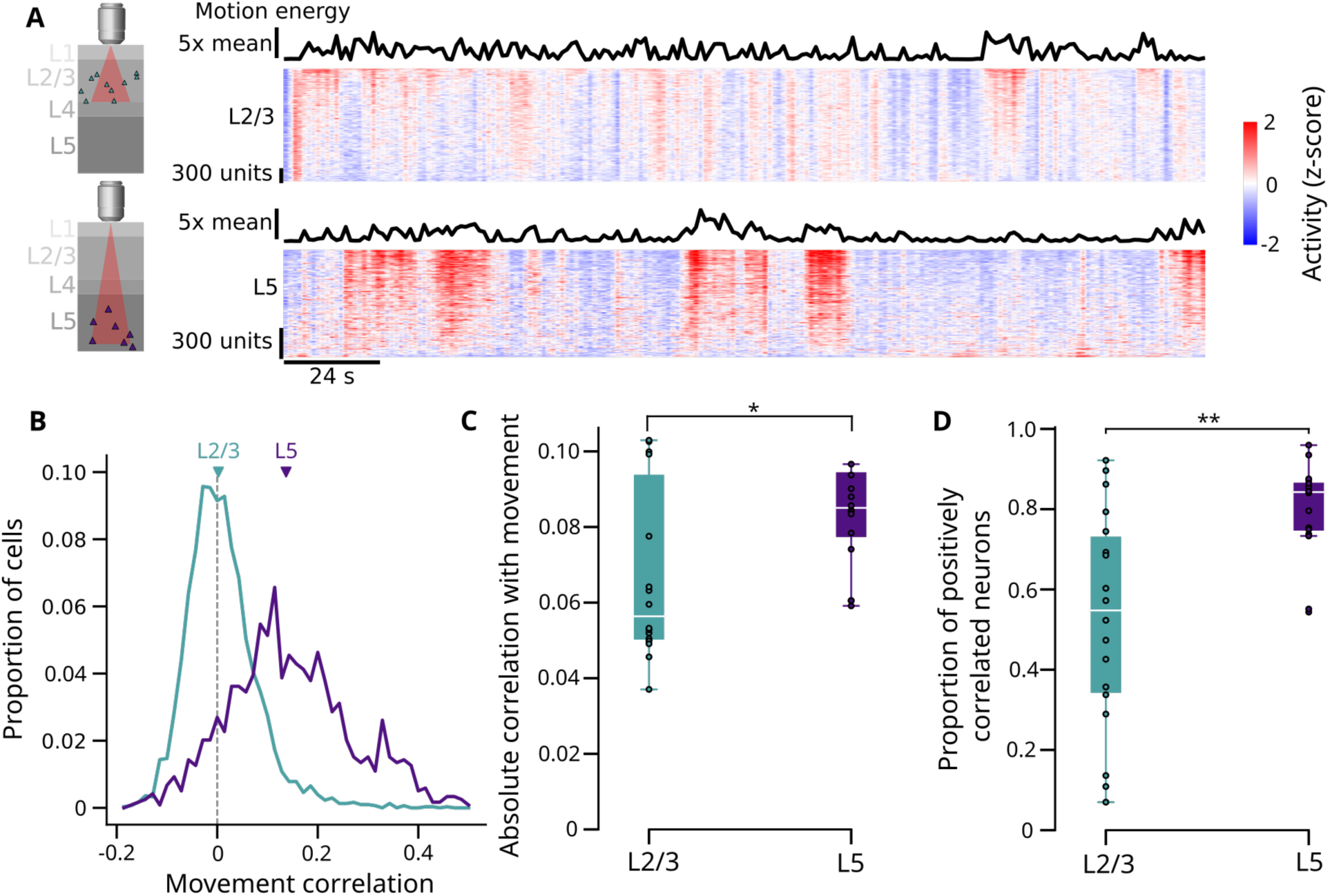
Movement correlates are more uniform in L5 than L2/3. (**A**) Top: example 2-photon recording of L2/3 spontaneous activity, with videographic motion energy shown as a black curve (sub-sampled to 2.5 Hz), and L2/3 population activity shown as a pseudocolor raster with cells vertically arranged by movement correlation (2.5 Hz sampling rate). Right: a curve showing the Pearson correlation of neural activity with movement energy (x-axis) for cells ordered as in the raster (y-axis). Bottom: same for L5 neurons. **(B)** Histograms of movement correlations of all cells in L2/3 (green) and L5 (purple) example experiments shown in (A). Triangles show mean movement correlation values of the example recording sessions. **(C)** Mean absolute correlation of neurons with movement, averaged over all cells in each session (one dot per session), in L2/3 and L5 (p = 0.02, rank sum test, N = 18 and 16 sessions in L2/3 and L5). **(D)** Proportion of cells in each session positively correlated with movement (p = 0.003), rank sum test, N = 18 and 16 sessions in L2/3 and L5). * p < 0.05, ** p < 0.01.

The different encoding of movement in L2/3 and L5 populations led to differences in movement decoding. We used linear regression to predict motion energy (i.e. the mean absolute derivative of face video pixel values) from the first 40 principal components of neural activity acquired using two-photon calcium imaging (Figure S4A, B). When decoding motion energy from the full population, performance was above chance from activity in either L2/3 and L5 (Figure S4C-E), however a difference became apparent when decoding from smaller population sizes, for which L5 predicted movement better than L2/3 (Figure S4G). This suggests that the stronger coding of movement information in L5 has led to a more robust code.

### Movement desynchronizes L2/3 but synchronizes L5

Movement decreased spontaneous population synchrony in L2/3 but increased it in L5. We assessed population synchrony by computing each neuron’s population coupling index^46^, defined as the Pearson correlation of its activity with the population rate (i.e. the summed activity of all recorded neurons). L2/3 excitatory neurons exhibited strongly synchronous spontaneous activity when the mice were stationary (Fig. 2A), which reduced when the animal moved, leading to a decrease in population coupling (Fig. 2B). By contrast, L5 spontaneous activity showed less synchronized fluctuations during stationary periods, but correlated better with running, leading to an increase in population coupling during movement (Figure 2C, D). The popula-tion coupling indices observed in L2/3 and L5 during both stationary and movement periods are significantly above chance level (p<.01; linear shift test; Figure S2A-B). This difference was consistent across sessions: for each cell we computed the ratio of population coupling coefficients between stationary and movement states (Figure 2E), and found that this ratio was significantly greater than 1 in L2/3 populations but significantly < 1 in L5 populations, with a significant difference between layers (Figure 2F; p =0.01, Linear mixed effects model; L2/3: N = 5 mice, 18 sessions; L5: N=5 mice, 16 sessions). Similar results were found with electrophysiology recordings (Figure S3E, F).

**Figure 2.**
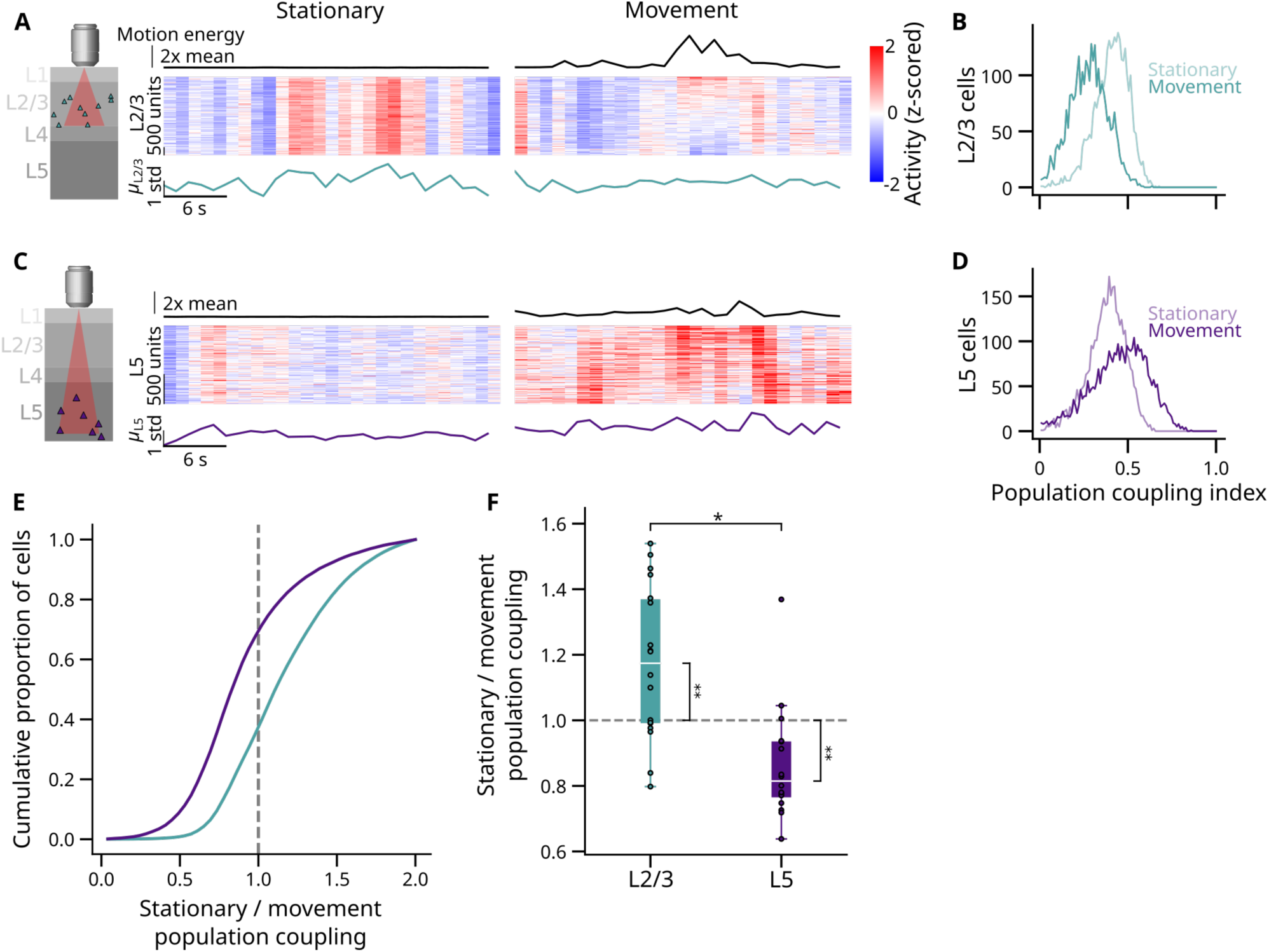
Movement desynchronizes L2/3 but synchronizes L5. **(A)** Pseudocolor rasters of L2/3 population activity during periods of station-arity (left) or movement (right), with neurons arranged vertically by correlation with movement as in Figure 1A. Above: videographic motion energy. Below, population rate, i.e. summed activity of all recorded neurons. **(B)** Histogram of population coupling indices (i.e. correlation of each neuron with population rate) for L2/3 neurons during movement and stationary periods. **(C-D)** Same as (A-B), for L5 neurons. **(E)** Cumu-lative histogram of the ratio of population coupling in stationary vs. movement periods, for L2/3 (green) and L5 (purple) neurons. **(F)** Comparison of the ratio of stationary and movement population coupling indices in L2/3 and L5 neurons. Each dot represents the mean coupling of all neurons in one recording session (p = 0.01, linear mixed effects model, N = 5 mice for both L2/3 and L5, N = 18 sessions for L2/3 and 16 sessions for L5). * p < 0.05, ** p < 0.01.

Thus, the most common causes of population synchrony are different between layers: In L5, spontaneous population synchrony mainly reflects a common positive modulation of firing by movement, while in L2/3, population synchrony instead primarily occurs due to internally-generated fluctuations that begin when movement stops.

### Spontaneous activity is lower-dimensional in L2/3 than L5

The strong population coupling seen in L2/3 spontaneous activity during stationarity suggests that the corresponding population activity may be dominated by a single dimension to which many neurons couple. To investigate this, we used shared variance component analysis (SVCA)^22^, which identifies activity dimensions which are reliably shared across cells and time, and quantifies the amount of neural activity reliably found in each dimension (Figure 3A, B).

**Figure 3.**
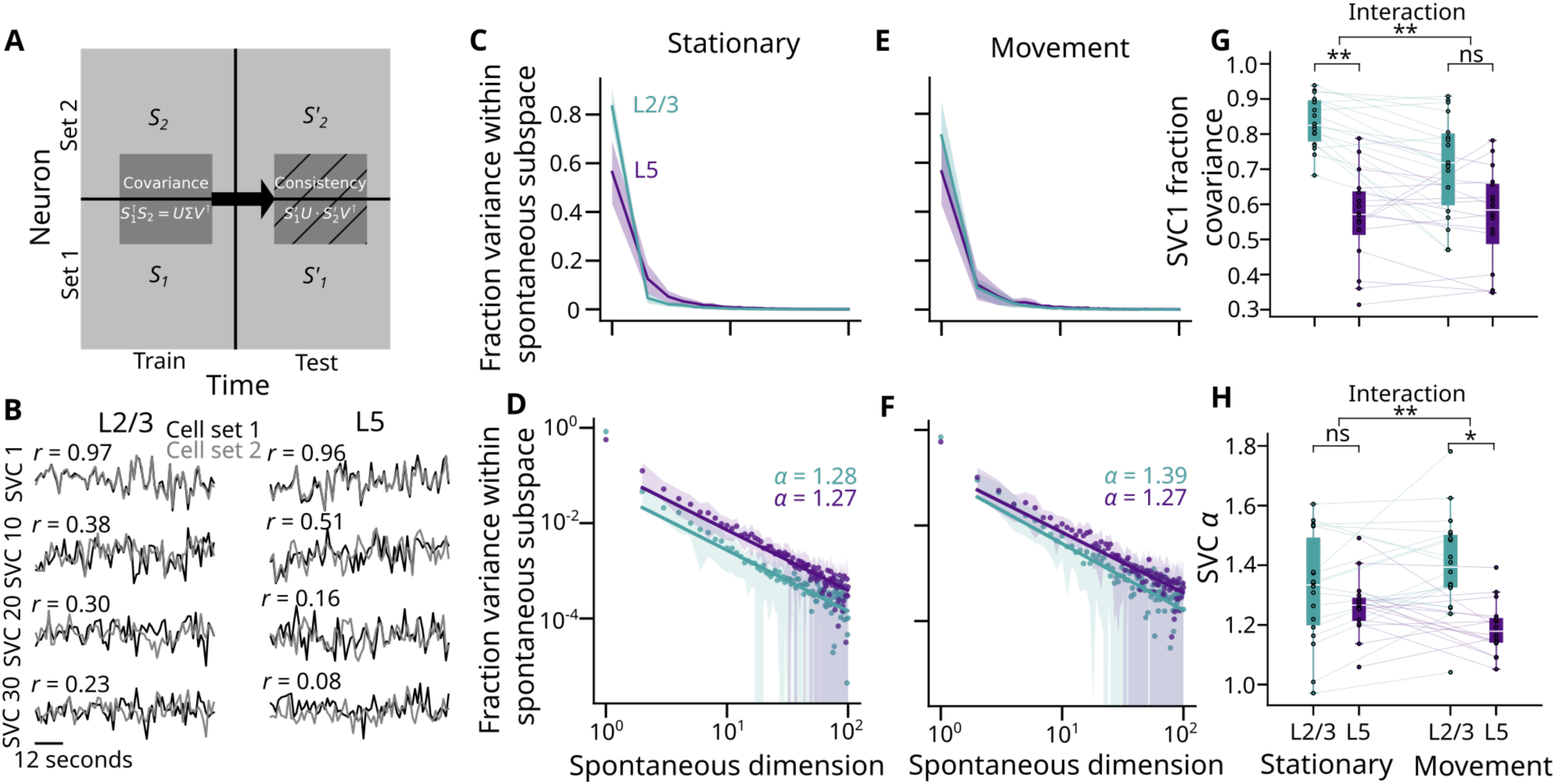
Spontaneous activity is lower-dimensional in L2/3 than L5. (**A**) Schematic of shared variance component analysis (SVCA). Neurons were randomly divided into two sets, and their covariance matrix was computed from training time points. The consistency of this shared activity was evaluated using testing time points. **(B)** Time traces of shared variance component projections in one example recording in L2/3 (left) and layer 5 (right), for 84 seconds of test-set activity. R-values represent the Pearson correlation between the shared variance component of cell set 1 and cell set 2 during testing time points. **(C)** Fraction of variance explained by each shared variance component in L2/3 (green) and L5 (purple) during stationary periods. Shading: standard error of the mean across mice. **(D)** As in (C), but with fraction variance in log-scale, and line of best fit of the second component onwards. **(E, F)** As in (C, D), but during movement periods. **(G)** Comparison of the fraction of variance explained by the first shared variance component in L2/3 and L5 populations during movement and stationary periods (p = 0.003 during stationary period, p = 0.15 during movement, interaction p = 0.003, linear mixed effects model. N = 5 mice for both L2/3 and L5, N = 18 sessions for L2/3 and 16 sessions for L5). **(H)** As in (G), but for the comparison of the variance-decay slope from the second component onwards (p = 0.570 during stationary period, p = 0.0212 during movement, interaction p = 0.001, linear mixed effects model). * p < 0.05, ** p < 0.01

SVCA analysis of an example session confirmed that spontaneous L2/3 population activity is more dominated by a single dimension than L5, particularly during stationary periods (for stationary periods the first dimension fraction variance for L2/3: 83% ± 2%, L5: 56% ± 3%, for movement periods: L2/3: 71% ± 3%, L5: 56% ± 3%, mean ± s.e.m, N = 18 sessions for L2/3 and N = 16 sessions for L5) (Figure 3C). Thus, the fraction of L2/3 variance from dimensions beyond the first was 1.7 times larger in running than stationary states (44% vs. 29%), and in stationary periods the fraction of variance from dimensions beyond the first was 2.6 times larger in L2/3 than in L5 (44% vs 17%). The reliable variance in all but the first few dimensions of spontaneous activity decays as a power law with exponent ∼1.3 (Figure 3D), as previously observed^22^. However, the first dimension had substantially higher variance than predicted by this power law, particularly in L2/3 in the stationary state. The first dimension accounted for significantly more variance in L2/3 than L5 during stationary, but not movement periods, and the interaction between layer and movement condition was significant (Figure 3G; Linear mixed effects model interaction term p = 0.003; N=34 sessions). The distribution of variance in dimensions beyond the first was less affected by movement, although the exponent of the power law was larger in L2/3 than L5, specifically during movement periods (Figure 3H).

Thus, L2/3 spontaneous activity is lower-dimensional than L5, and that the nature of this difference depends on state: during stationary states it is dominated by a single dimension, while in movement periods the power-law exponent is larger.

### Correlates of sound-evoked movements are larger in L5 than L2/3

We next investigated V1 responses to auditory stimuli in the different layers (Figure S5). Consistent with previous results^44^, auditory-evoked responses were strong when recorded with Neuropixels and correlated with sound-evoked movements, consistent with their interpretation as movement-related, rather than specifically auditory. These sound-evoked responses were stronger in L5 than L2/3, consistent with stronger movement correlates in deep layers (Figure 1). In 2p recordings, unlike Neuropixels recordings, sound presentation did not evoke body movements, because the background noise of the microscope prevents startle responses (Figure S5A-C). We did not observe auditory-evoked responses in V1 in 2p recordings, supporting the interpretation that sound presentation evokes V1 activity primarily due to evoked body movements, which were seen in Neuropixels but not 2p recordings.

### Visual responses are more dominant and stable in L2/3 than in L5

In contrast to movement correlates (which were stronger and more positive in L5), visual stimulus responses were stronger and more positive in L2/3 than L5. We evaluated each cell’s mean response to natural videos as Δ*F*/*F*_!_, where *F*_!_ is pre-stimulus baseline and Δ*F* is the average of activity during the movie minus pre-stimulus baseline. Responses were stronger in L2/3 compared to L5 (Figure 4A-D), both in terms of the absolute magnitude of response (Figure 4E) and the proportion of neurons with increased activity after visual onset (Figure 4F). Analysis of electrophysiology experiments produced consistent results (Figure S6A-C), and further suggested that the greater visual responses in L2/3 are primarily explained by the very low baseline activity levels in this layer (Figure S8G-I).

**Figure 4.**
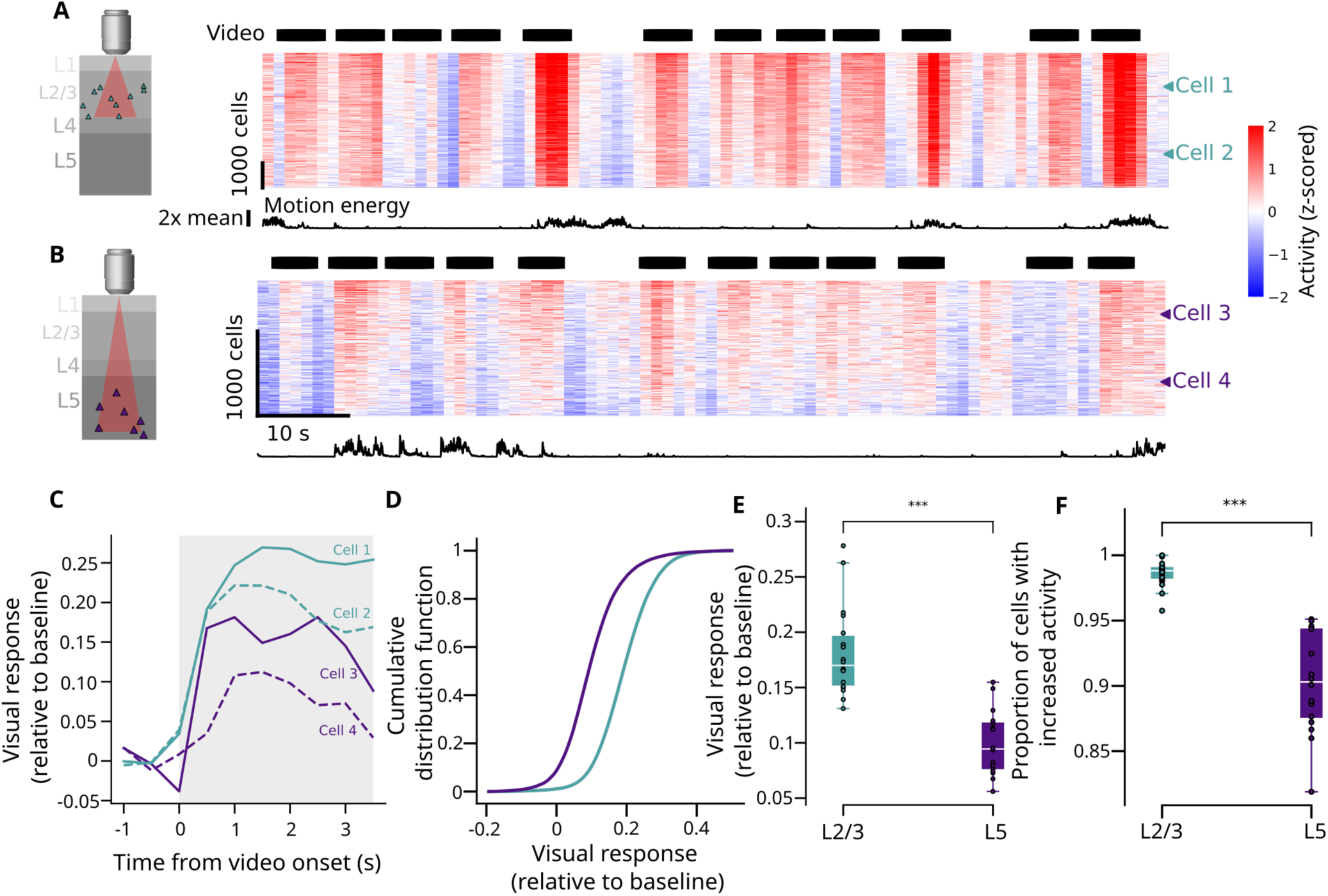
Visual onset responses are stronger in L2/3 than L5. **(A)** Pseudocolor activity raster for an example recording in L2/3. Black bars: times of video playback. **(B)** Same as (A) but for an example L5 recording. **(C)** Mean visual response across videos aligned to video onset (Z-scored relative to baseline) for four example cells indicated by arrows in (A) and (B). The gray region indicates the time window in which activity is averaged to compute video response shown in (D-F) **(D)** Histogram of mean visual response for cells in the example L2/3 (green) and L5 (purple) recordings. **(E)** Comparison of the mean absolute visual response between L2/3 and L5 neurons. Each dot represents the mean across neurons in a particular recording session (p < 0.001, rank sum test, N = 18 and 16 sessions in L2/3 and L5). **(F)** Same as (E), but for the proportion of neurons with increased activity after video onset (p < 0.001, rank sum test, N = 18 and 16 sessions in L2/3 and L5). * p < 0.05, ** p < 0.01, *** p < 0.001.

The results so far suggest that both L2/3 and L5 encode a mixture of visual and movement information, with a balance that differs between layers. To directly confirm this, we used a linear regression model to predict L2/3 and L5 activity from visual stimulus identity and video motion energy (Figure S9A). Movement features explained a greater proportion of population variance in L5 than in L2/3, and visual features explained a greater proportion of population variance in L2/3 than in L5 (Figure S9B). To further demonstrate the stronger encoding of movement activity in L5 than L2/3, we fit a visual encoding model using only trials where the animal was stationary, and evaluated this model’s performance on held-out stationary or movement trials (Figure S9C). We saw a reduction in explained variance when testing on movement trials in L5, but not L2/3 (Figure S9D, E), consistent with more purely visual code in L2/3. A visual decoding analysis showed consistent results: when decoding the visual stimulus from L5, but not L2/3 population activity, decoding performance was worse S10A-C). Again, this supports a more strongly visual code in L2/3 than L5.

### Stimulus-driven activity is lower-dimensional in L2/3 than in L5 and overlaps in only one dimension

To estimate the amount of stimulus-related power in successive dimensions we used cross-validated principal components analysis (cvPCA)^47^ (Figure 5A). This analysis was performed on data from movement and stationary states together, as it requires repeats of each stimulus which might have come in different states.

**Figure 5:**
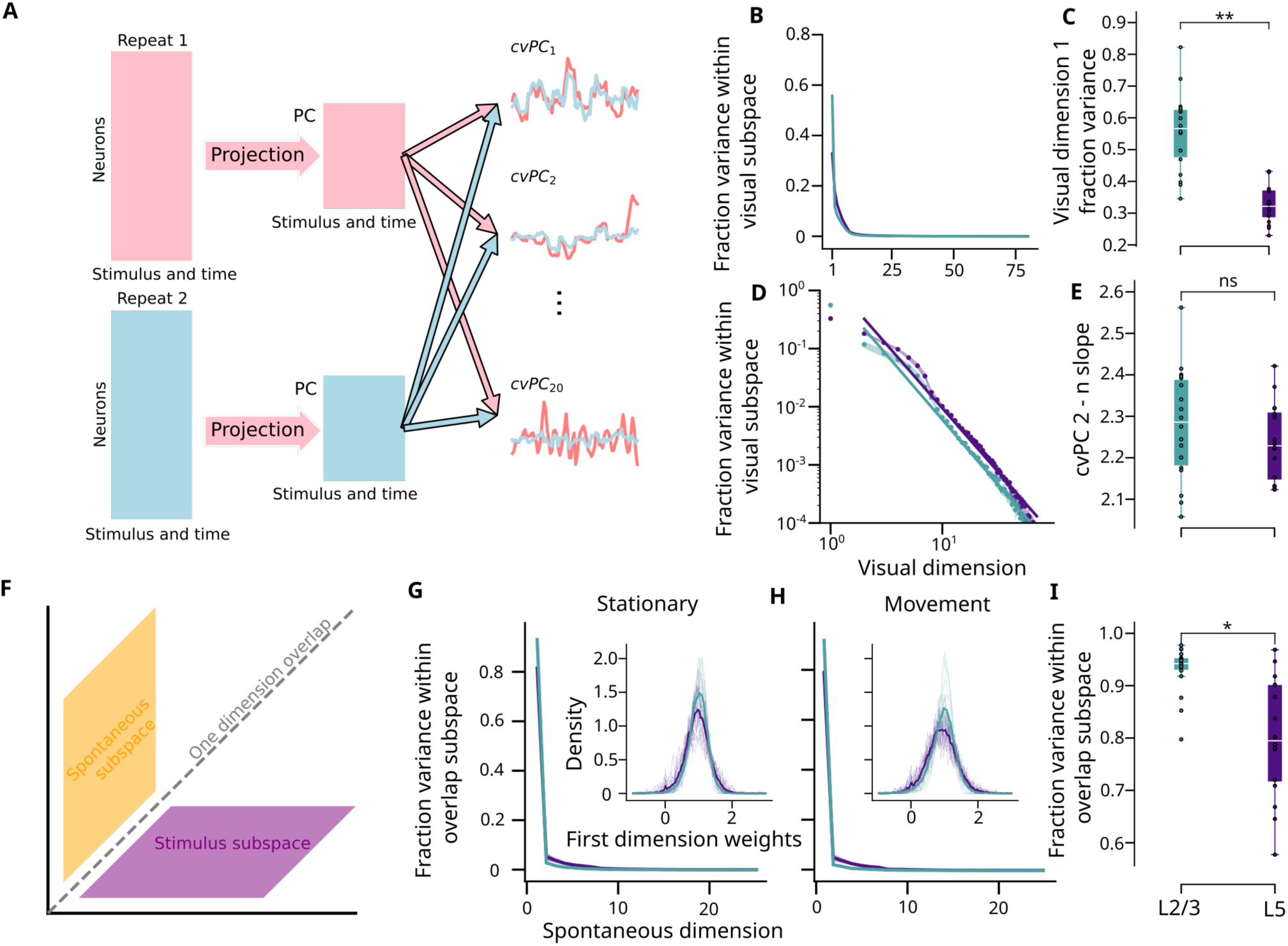
Population activity is more one-dimensional in L2/3 than L5. (**A**) Schematic of cross-validated principal components analysis (cvPCA). The matrix of stimulus-aligned responses across neurons in the first and second repeats are both projected using principal component weights learned from the first repeat. The covariance between the projected time courses of the two repeats provides an estimate of stimulus-related variance in each component. **(B)** Fraction covariance explained for each cross-validated principal component. (error bars showing stand-ard deviation too small to be visible). **(C)** Fraction of stimulus-related variance explained by the first visual dimension in L2/3 and L5 neurons. Each dot represents a recording session. (p = 0.007; N = 34 sessions, linear mixed effects model). **(D)** Same as (B), but with fraction variance on a log scale, and showing the line of best fit of variance decay from the second dimension onwards. **(E)** Same as (C) but comparing the variance decay slope between L2/3 and L5 neurons. **(F)** Schematic depicting the observation of a single dimension of overlap between the movement and visual subspace in the visual cortex. **(G)** Fraction of variance of the visual subspace that is explained by each spontaneous dimension, obtained during stationary periods. Inset: distribution of weights of individual neurons on the primary shared dimension. **(H)** Same as (G), but during movement periods. **(I)** Comparison of the fraction of variance in the first overlapping dimension between spontaneous and visual activity in L2/3 and L5 neurons. * p < 0.05, ** p < 0.01

The variances of visual responses were more dominated by the first dimension of activity in L2/3 than in L5 (Figure 5B-C). In L2/3, 56% ± 3% (mean ± s.e.m, N = 18 sessions) of reliable visual variance came from the first dimension, while in L5 33 ± 2% did (mean ± s.e.m, N = 16 sessions). Thus, the contribution of dimensions beyond the first was 1.5 times larger in L5 than in L23 (67% vs 44%). In higher dimensions, the geometry of population representations appeared similar: the slope of variance decay along the remaining dimensions was not significantly different between layers (Figure 5D-E).

The layers showed an even greater difference in the geometry of overlap between spontaneous and evoked activity. To estimate how the overlap between subspaces of spontaneous and evoked activity differed by layer, we projected visual activity onto the subspace defined by spontaneous activity, and then applied cvPCA to calculate the fraction of visual subspace activity that can be explained by spontaneous dimensions. Consistent with previous results^22^, we found that in L2/3 this overlap occurs almost exclusively in a single dimension (94% ± 1% of shared variance explained by this dimension, mean ± s.e.m, N = 18 sessions), and the weights of neurons onto this single dimension were nearly exclusively positive (Figure 5G-I). In L5 the largest shared dimension still had nearly all positive weights, but it accounted for a smaller fraction of the shared variance (80% ± 3%; Figure 5I, mean ± s.e.m, N = 16 sessions). Thus, the fraction of spontaneous-evoked overlap coming from dimensions beyond the first dimension is over three times larger in L5 than in L2/3 (20% vs 6%).

## Discussion

We found multiple differences between L2/3 and L5 visual cortex, in how visual and non-visual signals interact (Figure 6). First, activity in L2/3 is more strongly modulated by visual stimuli, while activity in L5 is more strongly modulated by movement. The way movement is encoded differs between layers: it increases the activity of nearly all neurons in L5, but in L2/3 an approximately equal fraction of neurons are excited as suppressed. Second, the structure of spontaneous activity in quiescent mice differs between layers. Quiescent-state spontaneous activity leads to synchronous fluctuations in both layers, but these are stronger in L2/3 and have a one-dimensional character. The overlap between spontaneous and evoked activity is also strongly 1-dimensional in layer 2/3, with <6% of the variance coming from dimensions beyond the first, in contrast to >20% in L5.

**Figure 6:**
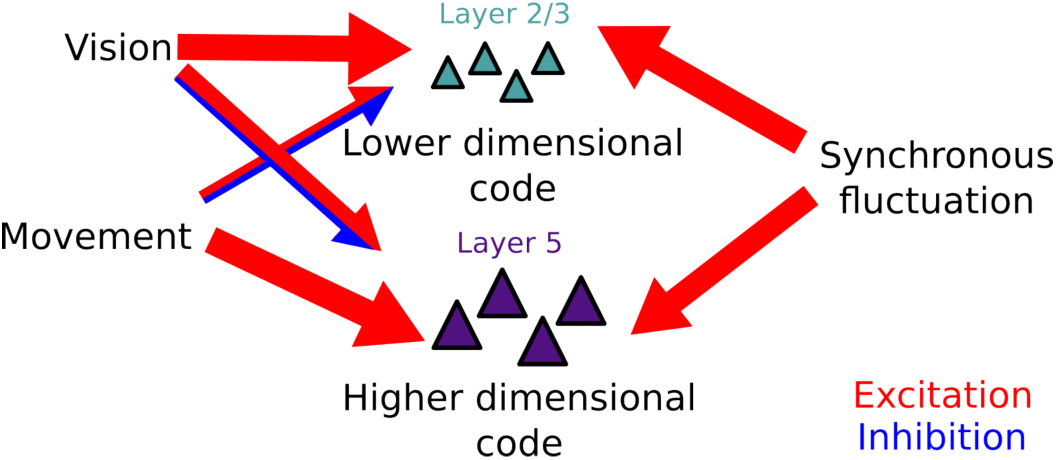
Summary of results. L2/3 population activity is lower-dimensional and more strongly modulated by visual stimuli and by resting-state synchronized fluctuations. L5 more strongly integrates information about bodily movement. Larger arrows indicate stronger modulation and red and blue components of an arrow indicate excitation and inhibition. These codes may be adapted for these layers’ synaptic targets in higher-order cortex and subcortical motor structures.

Both two-photon calcium imaging and extracellular electrophysiology are subject to experimental biases^33^. For imaging recordings, we controlled for differences in imaging quality between L2/3 and L5 cells (Figure S7), and for electrophysiology recordings, we controlled for differences in baseline activity (Figure S8). Our results were generally consistent between the two approaches, with one notable exception: auditory-evoked activity was found in all layers of visual cortex in electrophysiological, but not 2p recordings. We attribute this discrepancy to the suppression of sound-evoked behavioral responses (e.g. startle) by the background noise of the two-photon microscope, which was supported by abolition of startle responses when playing recorded 2-photon noise through a speaker in a test rig. Our data thus support the hypothesis that auditory responses in visual cortex arise primarily from stimulus-evoked movements^44^, and are thus stronger in L5 than L23 because L5 exhibits stronger movement correlates. On all other points, electrophysiology and 2-photon both showed similar effects, although the magnitudes of the effects sometimes differed. Specifically, electrophysiology showed that L2/3 neurons had larger baseline-relative sensory responses and synchronous-state fluctuations than L5 or L2/3 neurons, but the magnitude of this difference was larger when analyzed with 2-photon calcium imaging. We believe this occurs because electrophysiology spike sorting requires relatively high baseline firing rates^48^, which biases the result by preventing it from detecting sparsely-active excitatory neurons of L2/3 (Figure S8B).

The function of cortical spontaneous activity is one of the greatest mysteries of systems neuroscience. Our results do not resolve this question, but do provide important constraints on the answer. In particular, our results suggest that spontaneous activity should not be treated as a unitary phenomenon, but as containing at least two components, which are differently represented in different layers. The first component correlates with ongoing movements. This component is especially prominent in L5, where it positively modulates all neurons, and is weaker in L2/3, where it modulates around as many neurons positively as negatively. The second component consists of large spontaneous synchronized fluctuations that emerge during behavioral quiescence. This component modulates all neurons positively, and it is prominent but close to one-dimensional in L2/3, and weaker but higher-dimensional in L5.

One can imagine two functional interpretations for modulation of sensory cortical activity by movement. In the first interpretation, cortex combines multidimensional movement signals with multidimensional sensory signals to form a high-dimensional mixed representation of both streams^22,23^, which allows downstream structures to interpret sensory input in behavioral context in a flexible manner^25,26^. Under this hypothesis, our results suggest that the balance of sensory and movement information differs between layers: the cortical targets of L2/3 receive a greater balance of sensory information, while the often-subcortical targets of L5 receive a greater balance of motor information. This difference might be beneficial if subcortical structures use the information to produce motor outputs, for which ongoing motor context is more important, whereas higher-order cortex focuses on further sensory processing. The second interpretation for motor modulation of sensory cortex comes from the predictive coding hypothesis. In this view, movement signals in visual cortex represent predictions of sensory input due to self-motion, which can then be subtracted from actual sensory input^24,27–30^. Under this hypothesis, our results on the signs of modulation across layers are relevant. Our observation that negative modulation by movement is only seen in L2/3 is consistent with a hypothesis that L2/3 is specifically where subtraction occurs^27,29^; the fact that negative responses to sensory input occur primarily in L5 provides a further potential addition or correction to the framework.

For synchronous quiescent-state fluctuations, a first interpretation is that these reflect replay of prior sensory experiences, or hallmarks of an internal model of the world^11–16,31^. One of the main pieces of evidence for this view is the structural similarity of spontaneous and evoked population patterns. The low-dimensional nature of spontaneous activity in L2/3, and its even lower-dimensional overlap with sensory responses, therefore represents a challenge to the replay/model viewpoint, at least for superficial cortical layers. Importantly, even one-dimensional overlap can produce statistical signatures like those taken as evidence for internal models^32^, but one-dimensional overlap would be insufficient to genuinely model the sensory world. Thus, our results suggest that if the replay/model hypothesis does hold, it may hold specifically for deeper cortical layers. A second interpretation for quiescent-state fluctuations is that they represent “gating”^12,49^: an energy-saving mode in which cortex is only active part of the time, at the cost of slower processing speeds and reduced performance. Under this hypothesis, our data suggest that the cortex expends more energy to be active more often in L5 than in L2/3, perhaps because reduced quality output to L5’s subcortical targets carries a higher behavioral price than reduced quality output to L2/3’s cortical targets.

In summary, our data argue against a simple view in which all excitatory neurons within a cortical column participate in a common representation that is merely scaled across layers. Instead, superficial and deep excitatory populations in V1 differ systematically in the relative weighting of visual input, movement-related signals, and quiescent synchronized fluctuations. L2/3 carries a more visually dominated and more state-stable population code, while L5 carries a more movement-dependent code in which spontaneous and evoked activity overlap in a less exclusively one-dimensional manner. We therefore propose that the layered architecture of sensory cortex supports not a strict segregation, but a graded division of labour: superficial layers emphasize a robust sensory representation for downstream cortical processing, whereas deep layers more strongly combine sensory signals with behavioral context in a format suited to influencing subcortical and motor-related targets.

## Acknowledgements

We thank C. Reddy for support with surgery and lab management. M. Krumin for support with two-photon calcium imaging. R. Terry for animal breeding and maintenance. M.J. Wells for assistance in setting up the natural video protocol. A. Nunez-Elizalde for discussions during the conception of the natural video protocol. F. Takács and other members of the Cortexlab for discussions and comments. This work was supported by the Sainsbury Wellcome Centre Ph.D. program (T.P.H.S. and A.L.), Wellcome Trust (110120/185861 to P.C., 227065 to C.B., and 205093 and 204915 to M.C. and K.D.H.), and the EMBO (ALTF 740-2019 fellowship to C.B.). M.C. holds the GlaxoSmithKline/Fight for Sight Chair in Visual Neuroscience.

## Author contributions

T.P.H.S., P.C., K.D.H. and M.C. conceptualized the project and obtained the funds. T.P.H.S. performed the imaging experiments, C.B. and A.L. performed the electrophysiology experiments. T.P.H.S. performed the formal analysis and visualization with input from P.C. and K.D.H. T.S. wrote the original draft, and all authors contributed to the final version.

## Materials and Methods

### Surgery and recordings

Experimental procedures were conducted according to the UK Animals Scientific Procedures Act (1986) and under personal and project licenses released by the Home Office following appropriate ethics review.

For 2-photon calcium imaging, surgeries and recordings were performed on 10 mice, with 5 mice with the Ai94(tetO-GCaMP6s) x Camk2a-tTa x RBP4Cre genotype and 5 mice with the CamKII x tetO GCaMP6s genotype. Surgeries were performed under isoflurane anesthesia, and involved implantation of a headplate and a 3 or 4 mm circular craniotomy over the right primary visual cortex, as previously described^22,47,50^. Mice were habituated to handling and head-fixation before recordings were performed.

### Natural video stimulus presentation protocol

During each calcium imaging recording, mice were head-fixed, either with their forelimbs on a steering wheel^51,52^ (3/5 mice of each genotype) or on top of a treadmill (2/5 mice of each genotype). Visual stimuli were presented on three screens that cover 135 degrees in azimuth and 45 degrees in elevation, and auditory stimuli were presented using a single speaker (MF1 Multi-field magnetic speakers, Tucker-Davis Technologies). Audiovisual stimuli were presented as previously described^44^, consisting of combinations of seven 4-second natural scene videos and the corresponding seven sounds, including a grey screen and no sound condition, for a total of 8 × 8 = 64 combinations. These 64 audio-video combinations were presented 4 times each in blocks, with a different order in each block, with a 2 – 4 second inter-trial-interval between each audiovisual pair chosen randomly from a uniform distribution, and a 10 minute spontaneous period between each set of presentations. The electrophysiology recordings analyzed in this paper are the same as those analyzed in our previous study^44^. These recordings used an 11×11 combination of video and sounds, however for the analyses of the current study we subsampled only the 8 by 8 combination that matched that in imaging experiments. The electrophysiology protocol is described in full detail in our previous study^44^, and we used the same data described in that study for our electrophysiological results in this study. The videos presented during imaging were a subset of those presented in the previous study (Extended Figure 1B of ref 30), and include videos of an eagle, a frog, crows, a donkey, a seal, a bear, a cat and a dog.

### Movement quantification

We quantified behavioral movement by orofacial motion energy, which is better correlated with visual cortical activity than either running or pupil diameter^53^. Orofacial movements were monitored by infrared videography. An infrared LED illuminated the mouse and a camera with an infrared filter recorded at 30 frames per second. Videos were processed using Facemap^22,54^ to estimate motion energy in a region-of-interest covering the eye, whiskers and face. The motion energy trace was down-sampled to 2.5 Hz to perform correlation analysis with 2-photon calcium imaging data. To define stationary and movement periods, we divided the instantaneous motion energy by its mean, and time bins below / above the 60^th^ percentile of motion energy for each session were assigned to stationary / movement epochs.

### Two-photon calcium imaging of layer 2/3 and layer 5 neurons

Neural activity was measured with a commercial two-photon microscope (Bergamo II, Thorlabs Inc) controlled by ScanImage^55^, with a laser wavelength of 920 nm, at a scan rate of 2.5 Hz per plane across 12 planes spaced at 30 µm. Imaging was performed through either 3 or 4 mm cranial windows. After surgery, we performed widefield retinotopy mapping and used the sign field to ensure that all future 2-photon recordings were in V1. In all 2-photon experiments we performed recordings at two depth ranges: brain surface to 360 µm deep, and 180 to 540 µm below the brain surface, in alternating blocks in an A-B-A-B pattern with the depth of the first block balanced across sessions to eliminate potential biases due to imaging quality degradation or stimulus adaptation over the experiment. Both depth ranges used 12 planes at 30 µm spacing, scanned at 2.5 Hz per plane.

We obtained recordings of L2/3 or L5 excitatory populations by a combination of transgenic mouse lines and depth. L2/3 populations were recorded from CamKII x tetO GCaMP6s mice, which express GCaMP6s in all excitatory neurons ^40^, with only cells at depths 210 – 270 µm from the brain surface retained for analysis. Layer 5 populations were recorded from Ai94(tetO-GCaMP6s) x Camk2a-tTa x RBP4Cre mice ^41,42^ ^43^, which express GCaMP6s only in layer 5 excitatory neurons (both IT and ET cells), with only cells at depths 420 – 480 µm retained for analysis; although this depth range potentially overlaps with L4, L4 cells should not be labelled in this mouse line.

The laser power *P_z_* was shaped with depth *z* using the equation 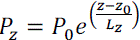, the percentage laser power at the most superficial depth *z*_0_, was set in the range of 10 – 30% for superficial imaging and above 30% for deep imaging, *z*_0_ is the depth of the most superficial imaging plane. The space constant *L_z_* was by default set to 180 µm, but adjusted in some recordings to 300 or 500 µm to prevent saturation of recorded signals. Power was capped at 100% if the formula exceeded this value.

### Extraction of pixel fluorescence signal from detected ROIs and corresponding neuropil region

To investigate and mitigate potential differences in recording quality with ROI size (which in turn can differ between layers), we investigated whether ROI size correlates with visual response reliability and noise level. We found a positive and negative correlation, respectively (Figure S7), indicating that without correction, we would find an erroneous bias towards more reliable, lower noise recordings in larger cells. To avoid this potential confound, we took the activity of each soma as the average fluorescence of the center 4 pixels of the region detected using Suite2p ^56^ (Figure S7).

### Noise estimation in two-photon calcium imaging recordings

To quantify the noise level in fluorescence obtained from individual pixels or the mean fluorescence across multiple pixels corresponding to a particular region of interest, we first computed the Δ*F*/*F* of the fluorescence signal:

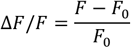

where *F* is the fluorescence signal and *F*_0_ is the mean fluorescence across time for the entire recording. We then computed the noise metric *ν* introduced in Ref. ^57^:

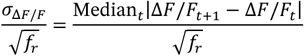

where *ν* is the estimated noise level, ΔF/F_t + 1_ − ΔF/F_t_ is the difference in signal between subsequent frames, and *f*_(_ is the frame rate of the recording (which is 2.5 Hz in all recordings in the present analysis). All analyses based on comparing properties of single cells (Figures 1, 4, S1, S4, S5G) only analyzed neurons with a noise level of 25% or below. Analyses of population geometry (Figures 2, 3,5) did not apply the threshold, to ensure larger population size.

### Estimating layer depth from Neuropixels recordings

We estimated laminar depth in Neuropixels recordings following Ref. ^58^, using the code at https://github.com/haiderlab/Current-Source-Density. Briefly, current source density analysis (CSD) was performed on the average local field potential aligned to video onset. The top of layer 4 was identified as the earliest and largest sink in CSD, and the depth of all units is defined relative to the top of layer 4.

### Separating visual and auditory-evoked components

To isolate the visual-related and auditory-related components of each cell’s fluorescence response we used an ANOVA-like decomposition ^44^. Each neuron’s response can be written as:

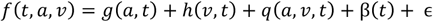

where *f*(*t*, *a*, *v*) is a cell’s signal as a function of time *t*, auditory stimulus *a* and visual stimulus *v*; *g*(*a*, *t*) is the auditory component of the cell’s response as a function of auditory stimulus and time; ℎ(*v*, *t*) is the visual component of the cell’s response as a function of the visual stimulus and time; *q*(*av*, *t*) is the interaction component of the cell’s response as a function of the audiovisual stimuli and time, β(*t*) is the mean component of the cell’s response independent of stimulus identity, and ϵ is the trial-to-trial variability that is not modelled.

To estimate the visual and auditory components of stimulus responses, we first z-scored each cell’s activity relative to the ensemble of pre-stimulus activity (all timepoints within 0-1 s prior of any trial) to yield an array *f*(*t*, *a*, *v*) for each cell, whose pre-stimulus activity had zero mean by definition. We then estimated the grand mean component of each cell’s response by taking the average activity over all videos and sounds:

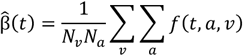

Where 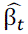 is the estimated mean response, *N_a_* is the number of auditory stimuli, and *N_v_* is the number of visual stimuli.

To obtain the visual-related component, we take the cell’s response over all possible sounds, and subtract that by the mean response component:

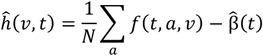

To obtain a single number quantifying how much each neuron was driven by auditory stimuli (Figure S5), z-scored 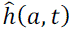 to an ensemble of that neuron’s activity in the range t= – 1 to 0 s prior to stimulus onset. We then take the average response timecourses across all sounds, to yield an auditory-evoked response timecourse for each unit over the range 0 – 4 seconds relative to stimulus onset and summarized this by averaging over time.

To quantify visual responses in imaging (Figure 4C-F) and ephys recordings (Figure S6A-C, S8G-H), we took trials where a video was presented, and computed the mean pre-stimulus activity from –1 s to 0 s prior to stimulus onset and the mean post-stimulus activity as 0 – 4 seconds after the stimulus onset. We then define the visual response relative to baseline as the post-stimulus activity subtracted by the pre-stimulus activity, and divided by the pre-stimulus activity. For ephys recordings, we also computed the mean post-stimulus activity during trials where a video was presented divided by the mean post-stimulus activity during trials where no videos were presented, and obtained similar results (Figure S8I).

To investigate the effect of time bins on the reliability of visual responses (Figure S8J), we quantified the stimulus-related variance of each neuron as:

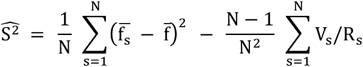

Where *N* is the number of distinct stimuli (treating each distinct time point in a video as a separate stimulus), *R_s_* is the number of repeats for stimulus *s*. *f_s,r_* is the response on repeat *r* to stimulus *s*. *f_s_* is the mean response to stimulus *s* across repeats, *f* is the grand mean across stimuli, and 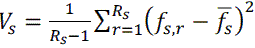 is the unbiased stimulus-independent variance for stimulus *s*.

### Linear shift test of movement correlation

To test for a significant correlation between the time courses of movement and neural activity (Figures S1, S4), we performed a linear shift test^45^. For each neuron, we computed the Pearson correlation coefficient r between its time course and the movement activity time course with shifts from –200 to 200 frames (1.2 seconds per frame). We then use a p-value threshold of 0.05 to indicate statistical significance, detecting a significant correlation if the unshifted r value is below the 2.5 or above the 97.5 percentile of the ensemble of all r values.

### Regression of motion SVDs using neural activity

To test how well neural activity in L2/3 and L5 predicts individual components of facial movement. We regressed motion SVDs from neural activity using two regression techniques; ridge regression and reduced rank regression. For each recording, we obtained the top 100 motion SVDs and subset only time periods where there is significant movement as previously described. Using neural activity with noise level below threshold (see “Noise estimation in two-photon calcium imaging recordings”), we fit a ridge regression model (α=10) and assessed quality of fit by performing 10-fold cross-validation and computing the Pearson’s correlation between the observed and predicted motion SVDs. We also fit a reduced rank regression model with rank 40, which predicts all 100 motion SVDs jointly. To ensure prediction performance is not due to differences in number of neurons per recording, we subsampled 150 neurons without replacement and repeated this procedure 5 times to obtain a mean.

### Shared variance components analysis

The quantification of overlap between spontaneous and visual subspaces was performed as in Ref. ^22^. To obtain estimates of the spontaneous variances we applied shared variance component analysis (SVCA) separately during stationary and movement periods (Figure 3). The fluorescence signal of each cell was first z-scored across the entire recording. We then partitioned the cells into a train and test set (with number of cells *N*_1_and *N*_2_) by dividing the imaging field-of-view into a 4 by 4 grid, and taking cells from alternating grid squares. We also partitioned time points into train and test set (with number of time points *T*_1_ and *T*_2_) by taking alternating 60 second segments. This yielded 4 activity matrices: 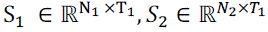 containing the activity of cell set 1 and 2 during the train set time points, and 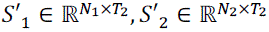 containing the activity during the test set time points. We then compute the covariance between cell set 1 and cell set 2 during the train time points and performed singular value decomposition with rank *r* on this covariance matrix to obtain our shared variance components (SVC) for cell set 1 in 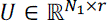 and cell set 2 in 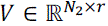

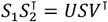

We then projected the corresponding cell set’s activity during the train time points on these shared variance components

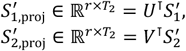

The shared variance along each SVC was estimated by the covariance between these two sets of projected cell activity during the train time points: 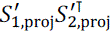

### Cross-validated principal components analysis (cvPCA)

To estimate the variance in successive dimensions of signal coding subspace we used cross-validated principal components analysis (cvPCA) ^47^.

We divided the repeats of each stimulus into two halves, and used each half *i* to obtain a matrix 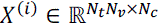 containing the concatenated timecourses of all stimulus responses, z-scored relative to pre-stimulus baseline activity as described above, for all cells (*N_t_* is the number of time points aligned to stimulus onset, *N_v_*. the number of videos, and *N_c_* the number of cells). We performed principal components analysis on the first activity matrix 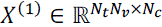 to obtain the principal components of visual activity, and projected the second-half of the data *X*^(2)^ onto these components, and calculated the covariance between the corresponding projections to obtain estimates of the visually-driven variances in the population (Figure 5B-E).

### Spontaneous and visual shared subspace analysis

To estimate the overlap between the spontaneous and visual subspaces (Figure 5G-I), we used the method of Ref. ^22^. Visual activity 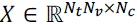 was projected onto the first 25 components of the spontaneous subspace identified with SVCA: 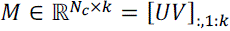, where *k* = 25, and cvPCA was performed on these projected components. The variance explained by each component quantifies the overlap between visual and spontaneous subspace. The weights of each neuron on the first shared dimension were obtained by calculating the linear combination of weights of the first shared dimension with the weights of each neuron on each of the first 25 spontaneous dimensions:

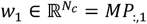

Where *P* ∈ ℝ*^k^*^×*p*^ is the loading of each the spontaneous subspace onto the principal components *p* obtained via cvPCA.

The weights were scaled for display to have a mean-squared value of 1 (Figure 5G-H, insets).

### Population coupling index

To quantify the synchrony of population activity, we computed the population coupling index of each neuron, defined as the Pearson’s correlation of the neuron’s activity with the mean population response:

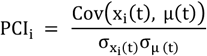

Where *x_i_*(*t*) is the activity of neuron i across time, and *μ*(*t*) is the mean activity of all neurons across time. To estimate the chance level values of population coupling indices, for each neuron, we performed linear shifts of its activity relative to the population across time, then re-calculated the population coupling indices during stationary and movement periods (Figure S2A), and computed the mean of these values across all shifts evaluated (Figure S2B).

### Relationship between firing rate and quantified metrics

To investigate the relationship between firing rate and the various metrics quantified in this study, we computed the number of spikes per second from our electrophysiology recordings in layer 2/3 and layer 5 units during the stationary period and movement periods (Figure S6A). We then took percentile bins of stationary firing rates (0-20%, 20 – 40%, 40 – 60%, 60 – 80%, 80 – 100%) for layer 2 / 3 and layer 5 units in each recording and visualised the absolute correlation with movement (Figure S8C), proportion of positively movement-correlated units (Figure S8D), visual response (Figure S8G), proportion of positively visual-modulated units (Figure S8H), and firing rate during visual stimuli divided by firing rate during absence of visual stimuli (Figure S8I).

### Regression models

To quantify the contributions of movement and visual stimuli to the responses in L2/3 and L5 cells, we fit 3 regression models to the neural activity recorded using imaging (Figure S9A-B). The first model consists of only the visual-stimuli regressors, which indicate the visual stimuli identity (which of the 7 videos) and have support from –1 to 4 seconds aligned to stimulus onset. The second model consists of only the movement regressors, which are the top 10 motion SVDs of the recording. The third model consists of both the visual-stimuli and movement regressors. In all three models, neural activity was binned at 1.2 s time bins, z-scored, then fitted using ridge regression with alpha = 10 and evaluated via 10-fold cross-validation.

To quantify whether there is a difference in variability of visual response due to movement between L2/3 and L5 neurons, we fit a visual-only model during stationary trials (defined as trials where the movement energy is below the 60th percentile), again using ridge regression with alpha = 10, and compared the explained variance when tested on the test set of stationary trials or movement trials (defined as trials where the movement energy is above the 60th percentile).

### Visual stimuli decoding

To compare the visual stimuli decoding performance of L2/3 and L5 cells within and across stationary and movement trials. We subsampled different number of neurons (2, 3, 4, 5, 10, 15, 20, 25, 50, 100, 150) and trained L2-regularised (with C = 10) SVM decoders to decode the stimulus identity from the population vector averaged from 0 – 4 seconds aligned to video onset. We computed the 2-fold cross-validated accuracy averaged across 200 subsample repeats for each neuron count. Stationary and movement trials were defined as before; based on whether the motion energy of each trial is below or above the 60th percentile. For visualization (Figure S10B), we performed linear discriminant analysis on 3 example videos in two recording sessions and plotted the projection of each trial onto the first two discriminant axes.

**Figure S1:**
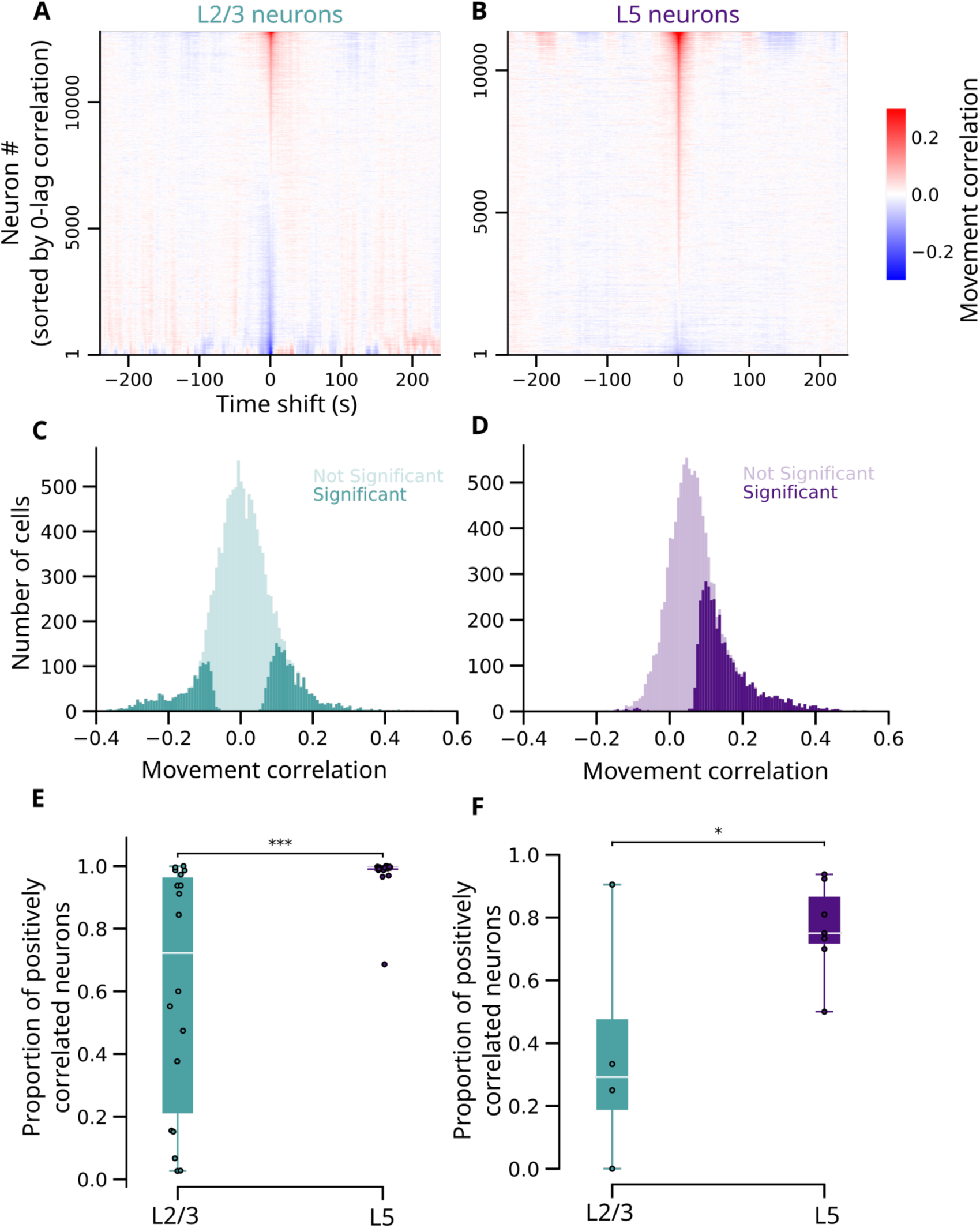
Testing neurons for significant movement correlation with the linear-shift test. **(A)** Heatmap of Pearson correlation values between fluorescence from neurons in L2/3 and lagged motion energy time courses, with different time shifts. Neurons are combined from all sessions and arranged vertically sorted by lag-0 correlation coefficient. **(B)** Same as (A) but for L5 neurons. **(C)** Histogram of correlation values in all recorded L2/3 neurons with significant (dark) and insignificant (light) correlation as assessed by the linear shift test. **(D)** Same as (C) but for layer 5 somas. **(E)** Proportions of significantly correlated neurons whose correlation is positive, in L2/3 and L5. Dots: individual recording sessions. The fraction in L2/3 and L5 are significantly different (p < 0.001, linear mixed effects model). **(F)** Same as (E), but for Neuropixels recordings (p < 0.05, linear mixed effects model).

**Figure S2:**
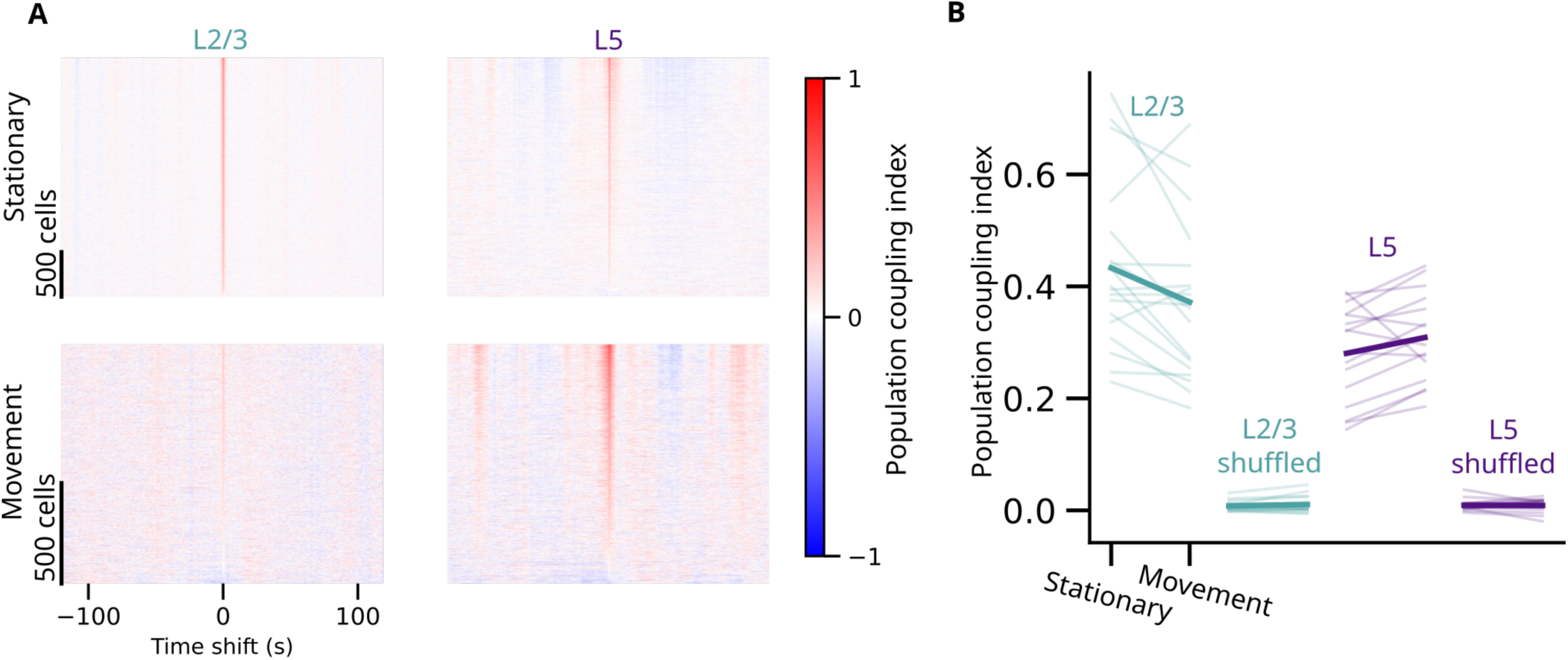
Testing neurons for significant movement correlation with the linear-shift test. **(A)** Heatmap of population coupling index of neurons in L2/3 (left) and L5 (right) during stationary (top) and movement (bottom) periods with different time shifts. Cells are sorted according to their 0-lag population coupling index. **(B)** Mean population coupling index for each recording session in L2/3 and L5 and the mean population index across time shifts (shuffles). The actual correlation was larger than all 99 shifts in each direction, giving significance at p=0.01.

**Figure S3:**
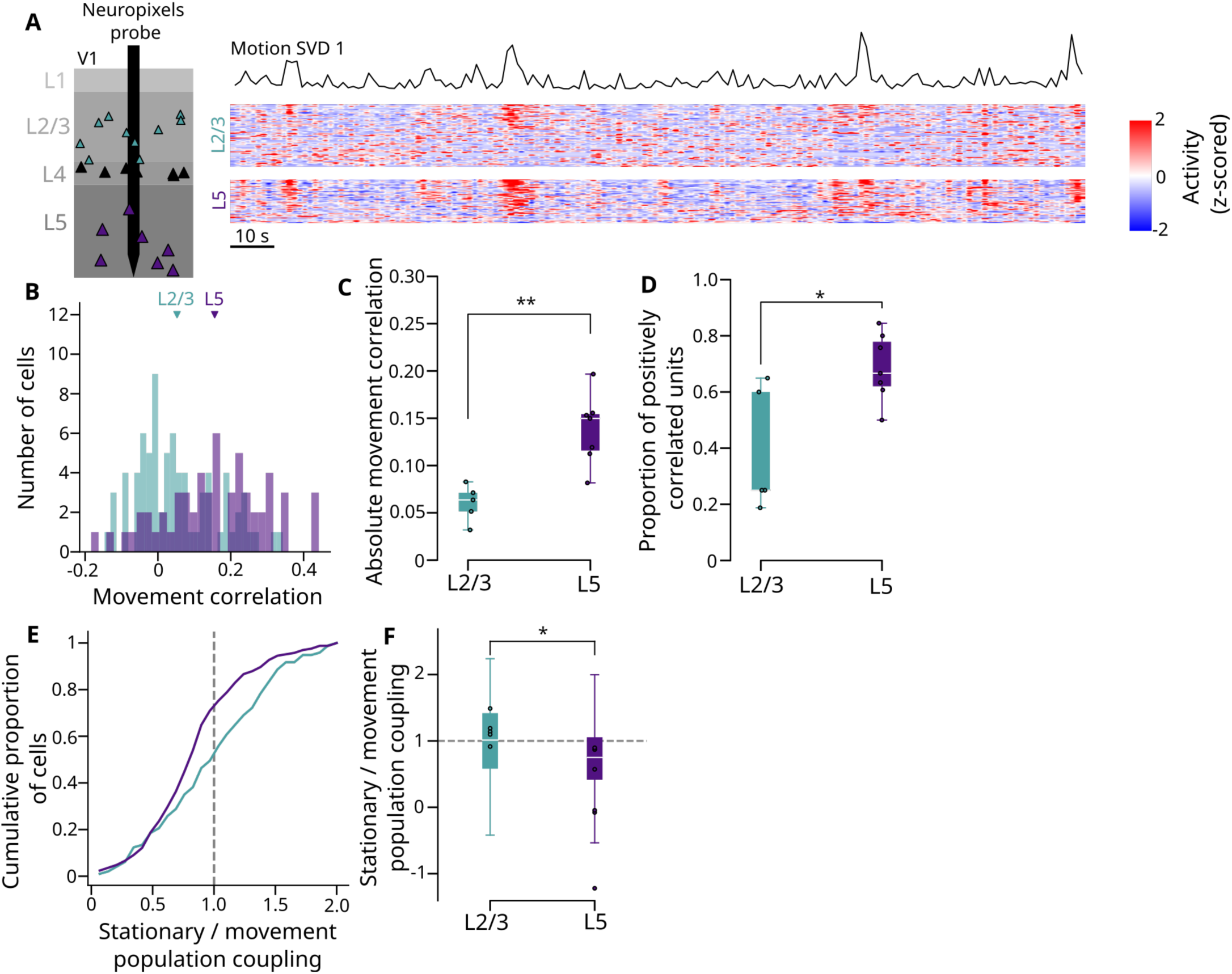
Movement correlation and population coupling in electrophysiology recordings. **(A)** Example recording using Neuropixels probes simultaneously in L2/3 and L5. Within each layer group, cells are sorted by signed movement correlation. **(B)** Histogram of signed movement correlation values of neurons in (A), including significant and non-significant correlations. **(C)** Comparison of the mean absolute movement correlation (significant or not) of neurons during each recording session in L2/3 and L5 (Difference between layers: p = 0.01, Wilcoxon rank-sum test, L2/3: N = 5 sessions; L5: N = 7 sessions). **(D)** Comparison of the proportion of cells positively correlated with movement (significantly or not) during each recording session (Difference between layers: p = 0.03, Wilcoxon rank-sum test, L2/3: N = 5 sessions; L5: N = 7 sessions). **(E)** Cumulative histogram of the ratio between the population-coupling index during stationary and movement periods across all recorded cells using Neuropixels. **(F)** Comparison of the population coupling index ratio in L2/3 and L5 (p < 0.01, linear mixed effects model). Note that electrophysiology recordings may under-sample sparsely firing neurons that may only fire during periods of strong network synchrony. * p < 0.05, ** p < 0.01

**Figure S4:**
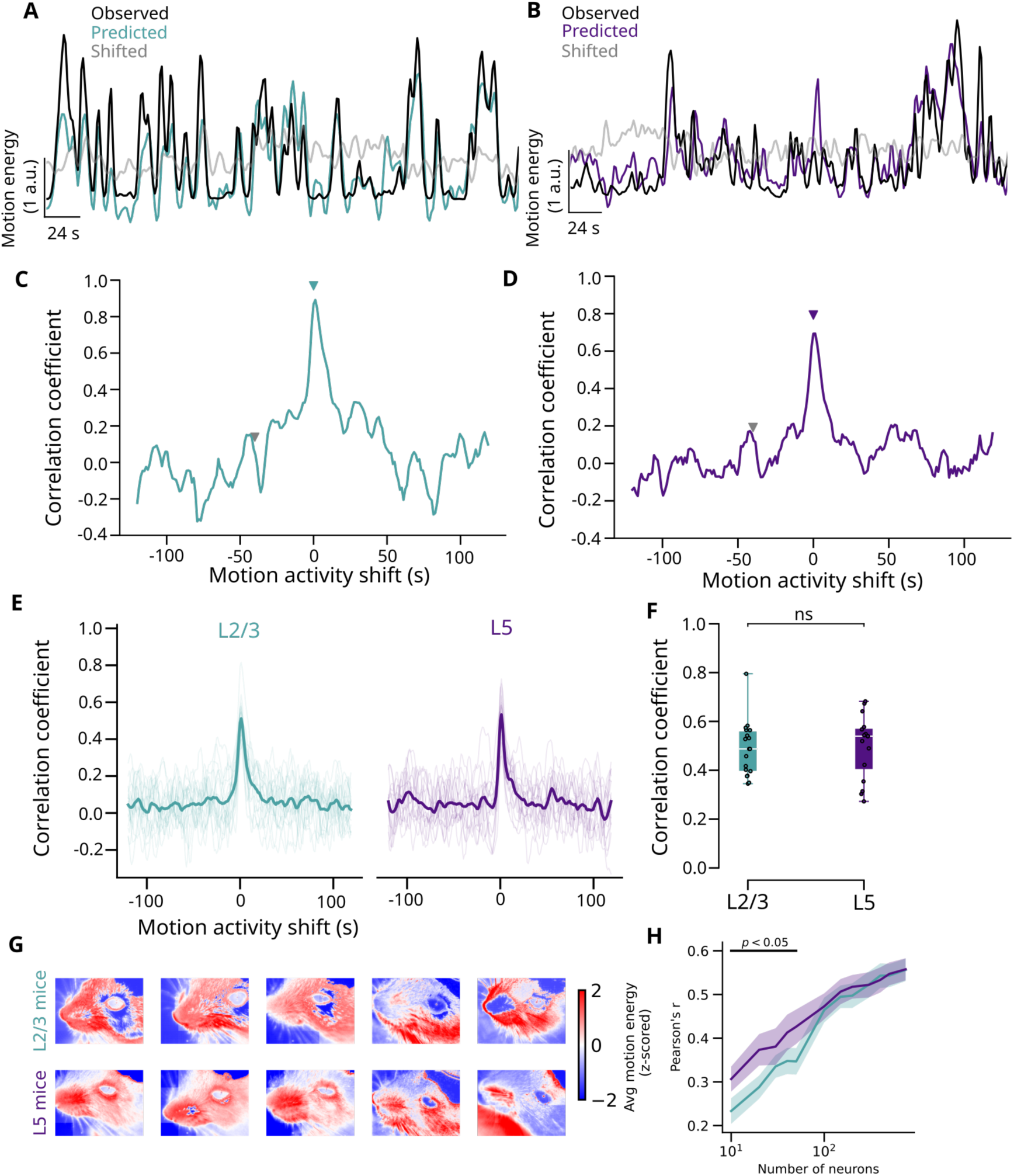
Regression and decoding of motion energy using neural activity. **(A)** Example motion energy trace (black) and motion energy predicted from L2/3 neural activity (green), for an example 2-photon recording. Grey line: prediction from neural activity time-shifted by –40 seconds (so neural activity lags). **(B)** Same as (A) but for an example 2-photon recording from L5. **(C,D)** Predictability of motion energy from neural activity, quantified by the Pearson correlation between observed and predicted motion energy, as a function of time shift, for these two example sessions. **(E)** As in (C-D), for all sessions (faint lines). Solid line: mean over sessions. **(F)** Comparison of zero time lag predictability for L2/3 and L5 (no significant difference between layers; *p* > 0.05), rank sum test across sessions. **(G)** Example motion energy maps of each mouse used in the study, i.e. the mean motion energy in each pixel, z-scored across pixels. **(H)** Pearson’s r between observed and predicted movement activity using different number of neurons as regressors (rank sum test). Significant differences in Pearson’s r was observed between 10 – 50 neurons.

**Figure S5:**
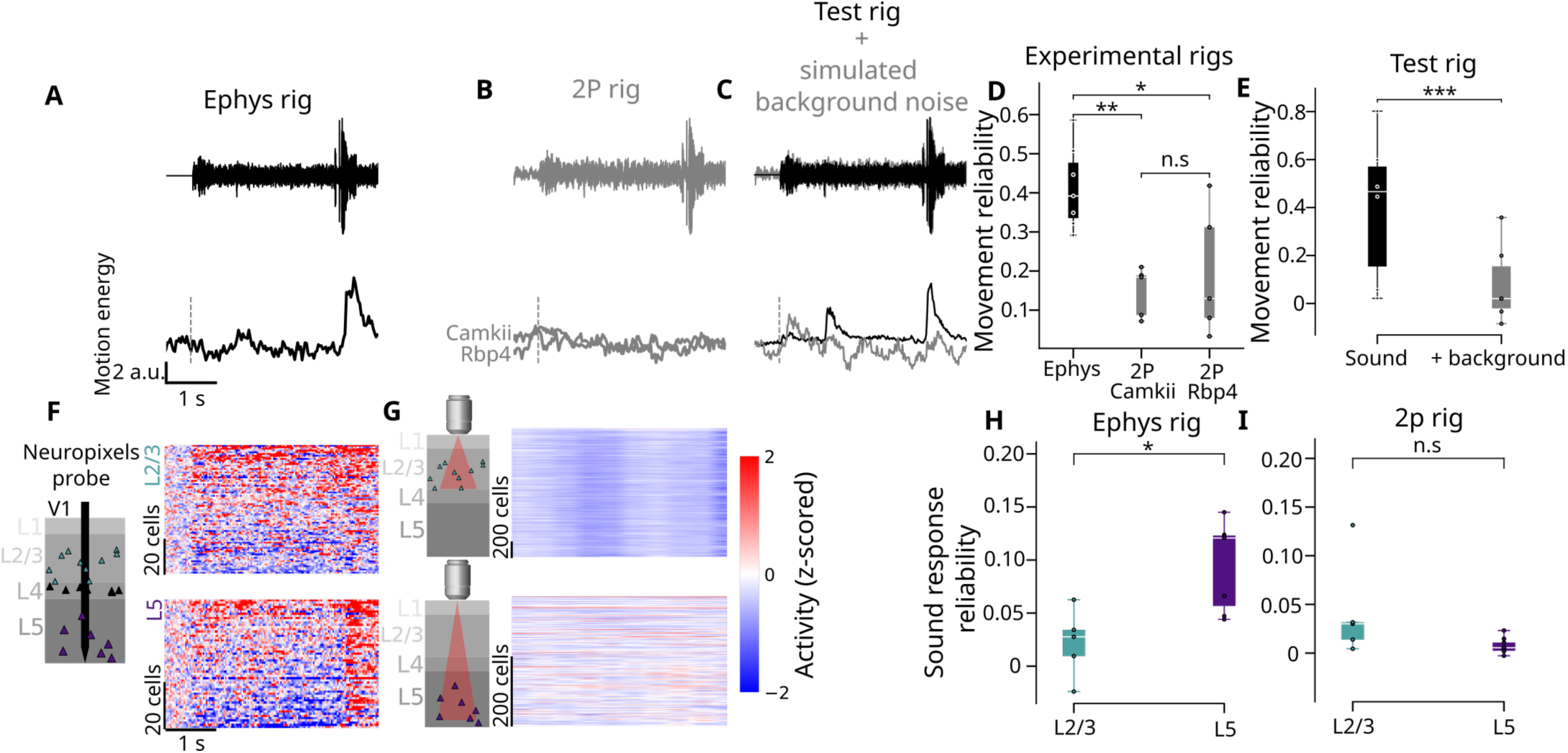
Differences in sound-evoked movement and neural response between electrophysiology and imaging recordings. **(A)** Top: envelope of an example sound played during electrophysiology recording. Bottom: corresponding motion energy trace averaged across trials from one example mouse. **(B)** Top: sound envelope including background noise in the two-photon imaging rig. Bottom: corresponding motion energy traces averaged over trials, from two example mice (one Camkii and one Rbp4). **(C)**. Top: sound envelopes representing the sounds played in the test rig, with and without simulated background noise. Bottom: trial-average motion energy traces with (gray) and without (black) background sound in one example session. **(D)** Boxplots comparing the reliability, quantified as the mean pairwise correlation of the movement traces across repeats, of auditory-evoked movements in mice recorded in the electrophysiology and imaging rigs. Sound-evoked movements were significantly more reliable in electrophysiology than 2-photon recordings (p = 0.004 between Ephys and 2P-Camkii, p = 0.042 between Ephys and 2P-Rbp4, p = 0.917 between 2P-Camkii and 2P-Rbp4, rank sums test across mice, N = 7 ephys mice, N = 5 for both Camkii and Rbp4 mice). **(E)** Boxplot comparing the reliability of auditory-evoked movements in the test rig with and without background sounds (p < 0.001, linear mixed effects model, N= 6 mice). **(F)** Pseudocolor raster for example L2/3 (top) and L5 (bottom) Neuropixels-recorded population, showing neural responses to the same sound as in (A) with aligned time axes. **(G)** Same as (F) but for example 2-photon session in L2/3 (top) and L5 (bottom), aligned to trace in (B). **(H)** Boxplot comparing the reliability of sound-evoked neural responses in layer 2/3 and layer 5 neurons acquired through electrophysiology (p < 0.05, rank sum test, N = 5 sessions in L2/3 and N=7 sessions in L5) **(I)** Same as (H) but for imaging recordings. (p > 0.05, rank sum test, N = 5 for both Camkii and Rbp4 mice) * p < 0.05, ** p < 0.01, *** p < 0.001

**Figure S6:**
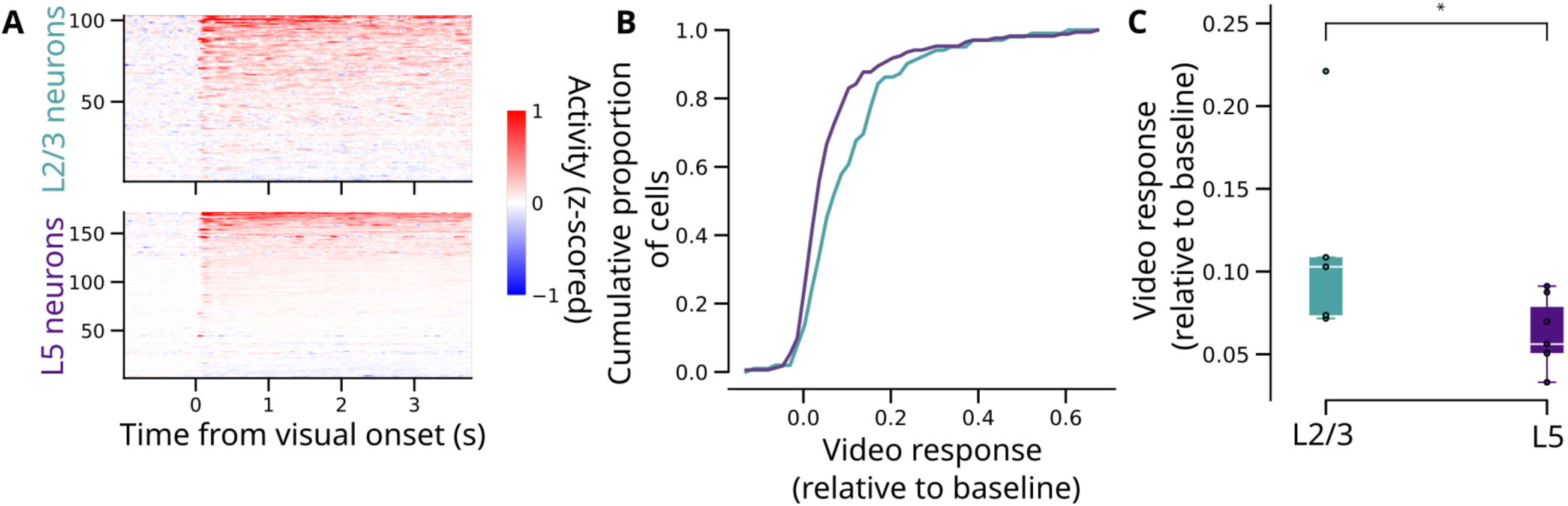
Visual responses in electrophysiology recordings. **(A)** Example recording using Neuropixels probes simultaneously in layer 2/3 and layer 5. Within each layer group, the cells are sorted vertically by their signed visual response. **(B)** Cumulative histogram of mean visual response of neurons in the example sessions of (A). **(C)** Comparison of the mean visual response of neurons in L2/3 and L5. Each point represents the mean visual response (mean across all sounds and from 0 – 4 s aligned to video onset) of all cells in a recording session. Visual responses are significantly larger in L2/3 than L5 (p < 0.05, linear mixed effects model, N = 5 sessions in L2/3 and N = 7 sessions in L5). * p < 0.05

**Figure S7:**
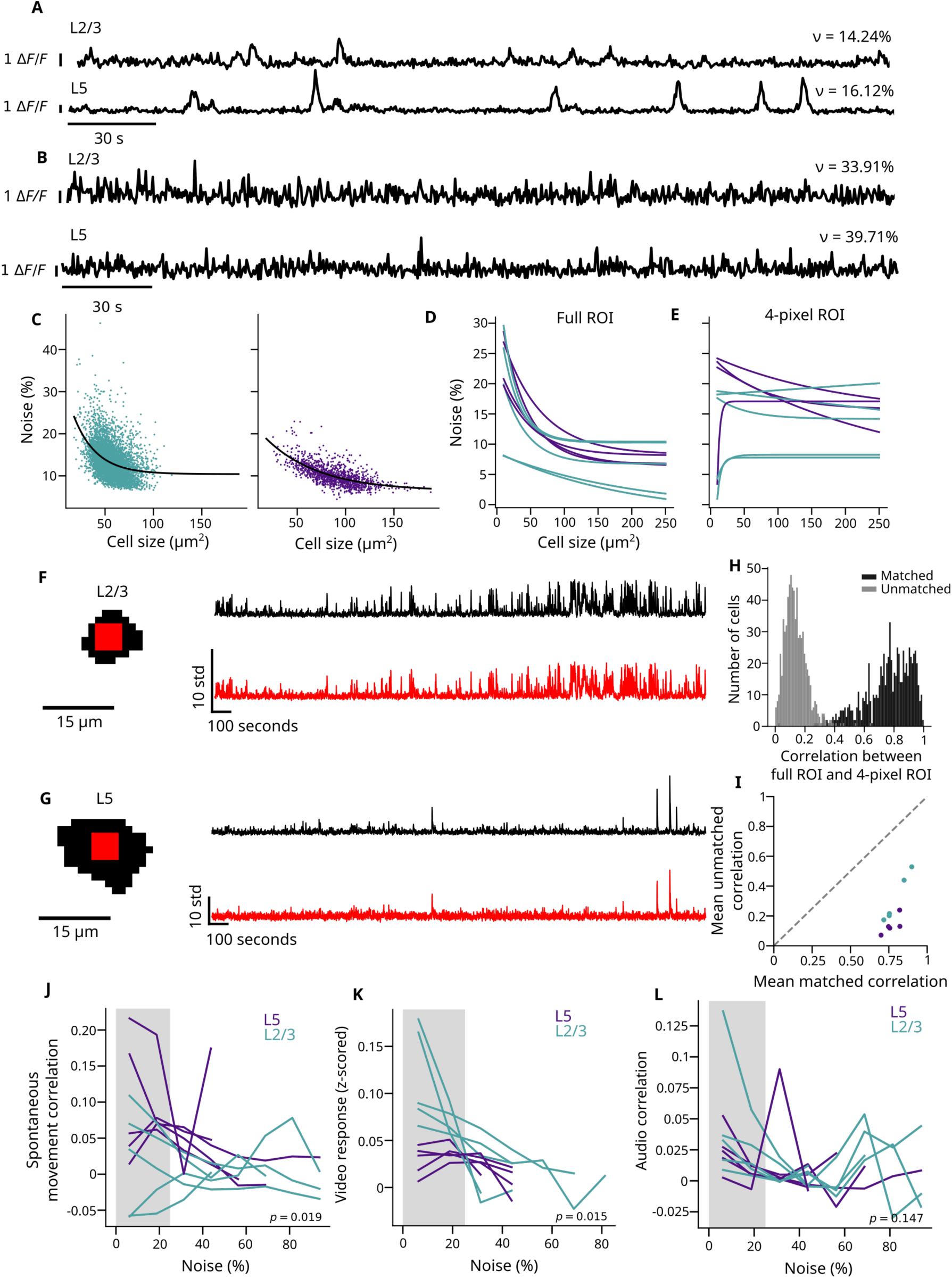
Controlling for the effect of soma size on noise levels. (**A-B**). Activity traces of two example L2/3 and L5 neurons with low and high estimated noise levels, together with estimates of noise metric *ν*. **(C)** Relationship between cell size and noise level of traces extracted from full ROIs for example sessions in L2/3 (left) and L5 (right). Black curve: exponential curves (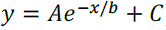) fit to these points by the Levenberg-Marquardt algorithm. **(D)** Superimposed curves showing exponential fits of relationship between cell size and noise level of traces extracted from full ROIs, for all sessions in L2/3 (green) and L5 (purple), each line represents the exponential decay curve fit to neurons obtained from each mouse. **(E)** Same as (D), but after controlling for size by using equally-sized ROIs consisting of the central 4 pixels from each ROI defined by suite2p. **(F)** Left: Silhouette of one example cell in L2/3 extracted using suite2p (black) and the center 4 pixels (red). Right: corresponding activity traces. **(G)** Same as (F) for an example neuron in L5. **(H)** Histogram over all ROIs of correlation coefficients between the full ROI activity and the 4-pixel ROIs activity (matched, black). Gray: histogram of full ROIs and 4-pixel ROIs of other units (cell identity shifted by 1; unmatched, gray). **(I)** Mean correlation between matched and unmatched full and 4-pixel ROIs for each mouse recorded in L2/3 (green) and L5 (purple). **(J)** Mean spontaneous movement correlation of each 4-pixel ROI in L2/3 (green) and L5 (purple) as a function of noise level. The gray region depicts the noise level range used in the cell-level comparisons in this study (Figure 1, 4, S5I). **(K)** Same as (J) but for video response (z-scored relative to baseline). **(L)** Same as (J) but for auditory response correlation.

**Figure S8:**
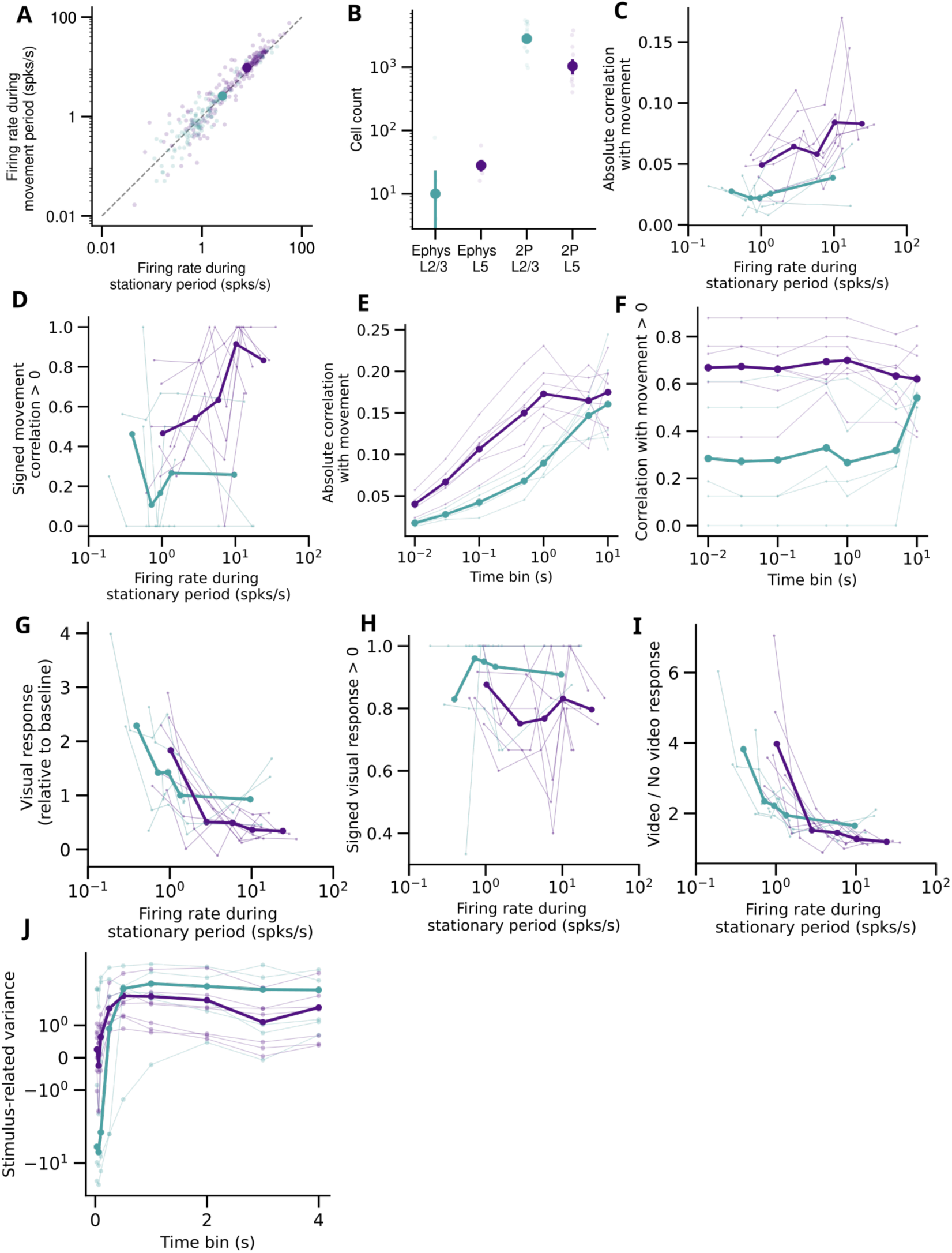
Relationship between absolute firing rate and quantified metrics, made using neuropixels data. **(A)** Spontaneous firing rate of L2/3 (green) and L5 (purple) units recorded using Neuropixels probes during stationary vs. movement periods (no visual stimuli). Small dots indicate individual units and big dots indicate the mean across all units. L5 firing rates are higher than L2/3 in both stationary periods (L23: 2.56, L5: 7.96 spikes/s, difference of 5.4 +/− 0.8, p < 0.01, linear mixed effects model) and movement periods (L23: 2.59, L5: 9.63 spike/s, difference of 7.0 +/− 0.98, p < 0.01, linear mixed effects model) **(B)** Cell count in electrophysiology and imaging recordings in L2/3 and L5. In ephys, more neurons were recorded from L5 (p < 0.01, paired t-test). In imaging experiments, more cells were recorded in L2/3 (p < 0.01, t-test). **(C)** The mean absolute correlation with movement as a function of firing rate of L2/3 and L5 units (percentile bins). Thin lines denote individual subjects and thick line denotes the mean across subjects. There is a significant main effect of both firing rate (p < 0.01) and layer differences (p < 0.001) on absolute movement correlation (linear mixed effects model). **(D)** Similar to (C) but for the proportion of units positively correlated with movement (p < 0.05 for firing rate main effect, p < 0.001 for layer main effect, linear mixed effects model). **(E)** The mean absolute correlation with movement across all L2/3 and L5 units as a function of the time bin used in binning neural activity and movement. Thin lines denote individual subjects and thick lines denote the mean across subjects. There is a significant main effect of both time bin (p < 0.001) and layer (p < 0.001) on movement correlation (linear mixed effects model). **(F)** Similar to (E) but for the proportion of units positively correlated with movement (p > 0.05 for time bin effect, p < 0.001 for layer main effect, linear mixed effects model). **(G)** Similar to (C) but for the visual response relative to baseline. There is a significant main effect of firing rate (p < 0.05), but not layer differences (p > 0.05) on visual response (linear mixed effects model). **(H)** Similar to (C) but for the proportion of units that are positively modulated by visual stimuli (p > 0.05 for firing rate main effect, p > 0.05 for layer differences main effect, linear mixed effects model). **(I)** Similar to (C) but for the firing rate during visual stimuli divided by the firing rate during blank periods (p < 0.01 for firing rate main effect, p > 0.05 for layer differences main effect, linear mixed effects model). **(J)** Stimulus-related variance in L2/3 and L5 neurons as a function of the time bin used to quantify stimulus response. Negative values indicate the stimulus-related variance is below noise level.

**Figure S9:**
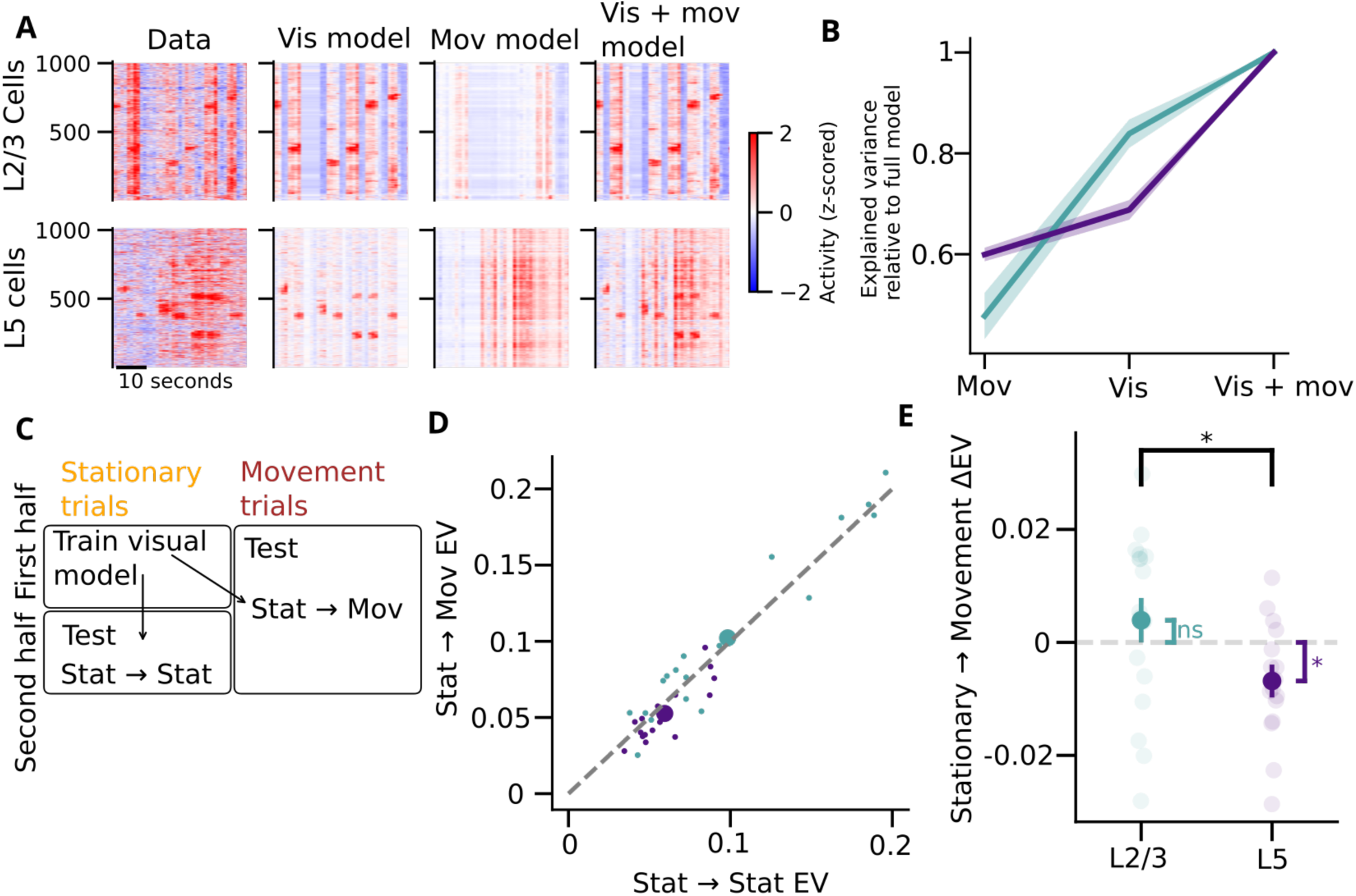
Regression of L2/3 and L5 response using movement and visual stimuli. **(A)** Example L2/3 and L5 rasters predicted by a regression model with visual stimuli, movement energy or both as regressors. **(B)** The explained variance relative to the full model (both regressors) for the movement-only model and visual-only model in L2/3 (green) and L5 (purple). Shaded region denotes the SEM across experiments. This analysis used all data (spontaneous and stimulus-evoked periods). **(C)** Schematic of a regression model with visual stimuli regressors that is trained during stationary trials, and tested on either another set of stationary trials or on movement trials. **(D)** Comparison of the explained variance of the visual model shown in (C) tested on stationary trials (x-axis) versus on movement trials (y-axis). Small dots denote individual experiments and big dots denote the mean across all experiments. This analysis used only the stimulus period (–1 to 4 s relative to the onset of the 4s movies). **(E)** Comparison of the difference between the explained variance by the visual model during movement and stationary trials. In L2/3 there is no significant difference (p > 0.05), and in L5 the explained variance is lower when tested on movement trials. * p < 0.05

**Figure S10:**
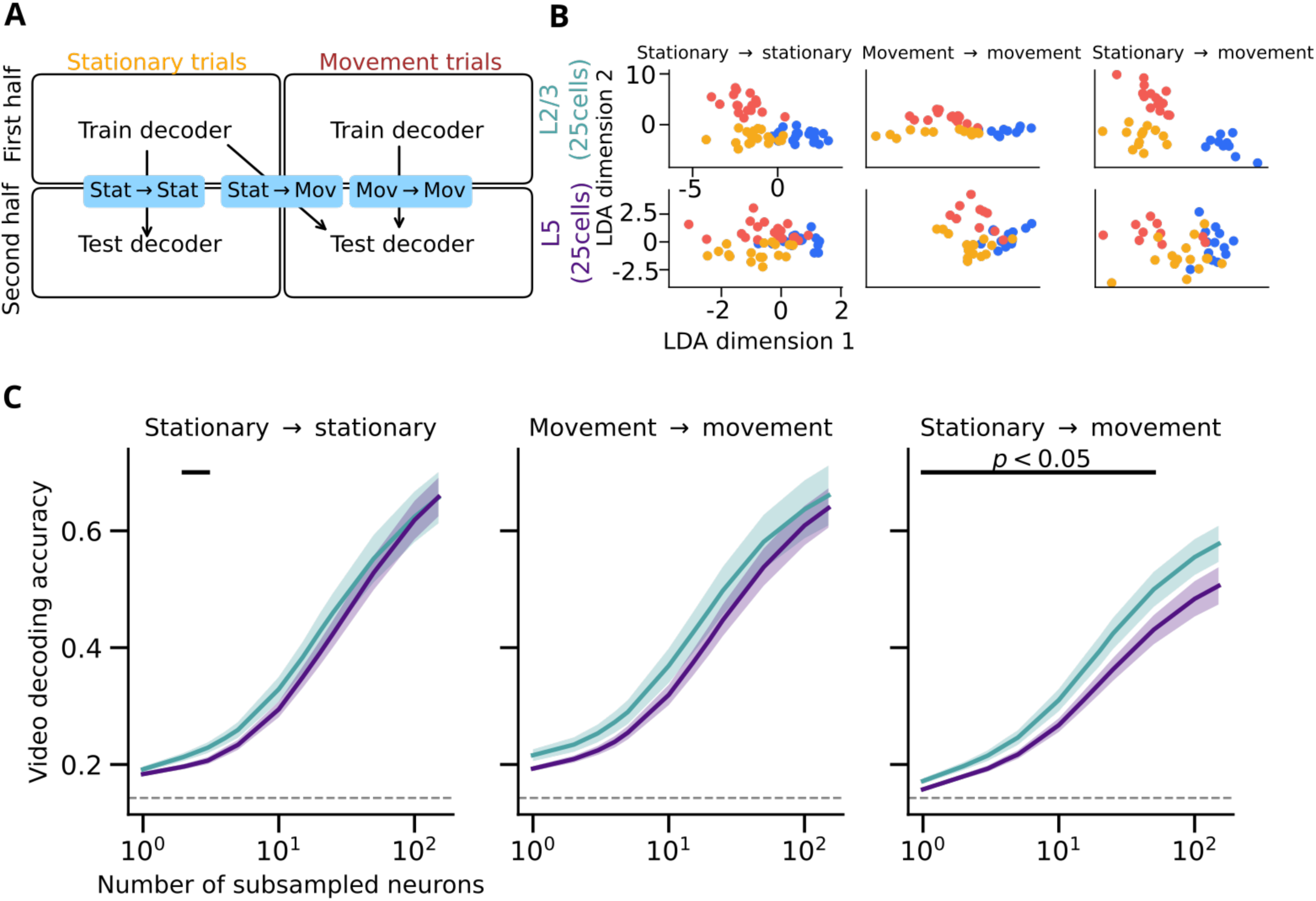
Decoding of visual stimuli across stationary and movement trials. **(A)** Schematic of visual stimulus decoder trained on either stationary or movement trials and tested on either stationary or movement trials. **(B)** Example of visual stimuli projected on the linear discriminant analysis decoding axes in L2/3 (top) and L5 (bottom) for each possible assignment of stationary or movement conditions to training and test sets (title: training set –> test set). Each point represents the mean response over the 4s presentation of one trial of one of three videos (video identity indicated by dot color). Note the greater separate in L2/3, particularly in the stationary->movement cross-decoding condition. **(C)** for each train-test condition, as a function of neuronal population size. Shading: standard error across experiments. Black bars: significant difference in decoding between L2/3 and L5. Dashed lines indicate chance decoding accuracy.

## References

1. Ringach, D. L. Spontaneous and driven cortical activity: implications for computation. Current Opinion in Neurobiology vol. 19 439–444 Preprint at 10.1016/j.conb.2009.07.005 (2009).

2. Arieli, A., Sterkin, A., Grinvald, A. & Aertsen, A. Dynamics of ongoing activity: Explanation of the large variability in evoked cortical responses. Science (1979). 273, 1868–1871 (1996).

3. Kenet, T., Bibitchkov, D., Tsodyks, M., Grinvald, A. & Arieli, A. Spontaneously emerging cortical representations of visual attributes. Nature 425, 954–956 (2003).

4. Han, F., Caporale, N. & Dan, Y. Reverberation of Recent Visual Experience in Spontaneous Cortical Waves. Neuron 60, 321–327 (2008).

5. Mohajerani, M. H. et al. Spontaneous cortical activity alternates between motifs defined by regional axonal projections. Nature Neuroscience 2013 16:10 16, 1426–1435 (2013).

6. Hoffman, K. L. & McNaughton, B. L. Coordinated reactivation of distributed memory traces in primate neo-cortex. Science (1979). 297, 2070–2073 (2002).

7. Ji, D. & Wilson, M. A. Coordinated memory replay in the visual cortex and hippocampus during sleep. Nature Neuroscience 2007 10:1 10, 100–107 (2006).

8. Fiser, J., Chiu, C. & Weliky, M. Small modulation of ongoing cortical dynamics by sensory input during nat-ural vision. Nature 2004 431:7008 431, 573–578 (2004).

9. Berkes, P., Orbán, G., Lengyel, M. & Fiser, J. Spontaneous cortical activity reveals hallmarks of an optimal internal model of the environment. Science (1979). 331, 83–87 (2011).

10. Luczak, A., Barthó, P. & Harris, K. D. Spontaneous Events Outline the Realm of Possible Sensory Responses in Neocortical Populations. Neuron 62, 413–425 (2009).

11. O’Neill, J., Pleydell-Bouverie, B., Dupret, D. & Csicsvari, J. Play it again: reactivation of waking experience and memory. Trends Neurosci. 33, 220–229 (2010).

12. Luczak, A., McNaughton, B. L. & Harris, K. D. Packet-based communication in the cortex. Nature Reviews Neuroscience vol. 16 745–755 Preprint at 10.1038/nrn4026 (2015).

13. Fiser, J., Berkes, P., Orbán, G. & Lengyel, M. Statistically optimal perception and learning: from behavior to neural representations. Trends Cogn. Sci. 14, 119–130 (2010).

14. Orbán, G., Berkes, P., Fiser, J. & Lengyel, M. Neural Variability and Sampling-Based Probabilistic Represen-tations in the Visual Cortex. Neuron 92, 530–543 (2016).

15. Haefner, R. M., Berkes, P. & Fiser, J. Perceptual Decision-Making as Probabilistic Inference by Neural Sam-pling. Neuron 90, 649–660 (2016).

16. Aitchison, L. & Lengyel, M. With or without you: predictive coding and Bayesian inference in the brain. Curr. Opin. Neurobiol. 46, 219–227 (2017).

17. Poulet, J. F. A. & Petersen, C. C. H. Internal brain state regulates membrane potential synchrony in barrel cortex of behaving mice. Nature 454, 881–885 (2008).

18. Niell, C. M. & Stryker, M. P. Modulation of Visual Responses by Behavioral State in Mouse Visual Cortex. Neuron 65, 472–479 (2010).

19. De Kock, C. P. J. & Sakmann, B. Spiking in primary somatosensory cortex during natural whisking in awake head-restrained rats is cell-type specific. Proc. Natl. Acad. Sci. U. S. A. 106, 16446–16450 (2009).

20. Schneider, D. M., Nelson, A. & Mooney, R. A synaptic and circuit basis for corollary discharge in the auditory cortex. Nature 513, 189–194 (2014).

21. Vinck, M., Batista-Brito, R., Knoblich, U. & Cardin, J. A. Arousal and Locomotion Make Distinct Contribu-tions to Cortical Activity Patterns and Visual Encoding. Neuron 86, 740–754 (2015).

22. Stringer, C. et al. Spontaneous behaviors drive multidimensional, brainwide activity. Science (1979). 364, (2019).

23. Musall, S., Kaufman, M. T., Juavinett, A. L., Gluf, S. & Churchland, A. K. Single-trial neural dynamics are dominated by richly varied movements. Nat. Neurosci. 22, 1677–1686 (2019).

24. Schneider, D. M. Reflections of action in sensory cortex. Curr. Opin. Neurobiol. 64, 53–59 (2020).

25. Fusi, S., Miller, E. K. & Rigotti, M. Why neurons mix: High dimensionality for higher cognition. Curr. Opin. Neurobiol. 37, 66–74 (2016).

26. Tye, K. M. et al. Mixed selectivity: Cellular computations for complexity. Neuron 112, 2289–2303 (2024).

27. Bastos, A. M. et al. Canonical microcircuits for predictive coding. [Neuron. 2012] – PubMed – NCBI. Neuron 2 76, (2012).

28. Audette, N. J. & Schneider, D. M. Stimulus-Specific Prediction Error Neurons in Mouse Auditory Cortex. Journal of Neuroscience 43, 7119–7129 (2023).

29. Keller, G. B. & Mrsic-Flogel, T. D. Predictive Processing: A Canonical Cortical Computation. Neuron 100, 424–435 (2018).

30. Miyashita, Y. Cortical Layer-Dependent Signaling in Cognition: Three Computational Modes of the Canonical Circuit. Annu. Rev. Neurosci. 47, 211–234 (2024).

31. Dimakou, A., Pezzulo, G., Zangrossi, A. & Corbetta, M. The predictive nature of spontaneous brain activity across scales and species. Neuron 113, 1310–1332 (2025).

32. Okun, M. et al. Population rate dynamics and multineuron firing patterns in sensory cortex. J. Neurosci. 32, 17108–17119 (2012).

33. Harris, K. D., Quiroga, R. Q., Freeman, J. & Smith, S. L. Improving data quality in neuronal population re-cordings. Nature Neuroscience vol. 19 1165–1174 Preprint at 10.1038/nn.4365 (2016).

34. Harris, K. D. & Thiele, A. Cortical state and attention. Nature Reviews Neuroscience vol. 12 509–523 Preprint at 10.1038/nrn3084 (2011).

35. Sakata, S. & Harris, K. D. Laminar-dependent effects of cortical state on auditory cortical spontaneous activity. Front. Neural Circuits https://doi.org/10.3389/fncir.2012.00109 (2012) doi:10.3389/fncir.2012.00109.

36. Bouvier, G., Senzai, Y. & Scanziani, M. Head Movements Control the Activity of Primary Visual Cortex in a Luminance-Dependent Manner. Neuron 108, (2020).

37. Flossmann, T. & Rochefort, N. L. Spatial navigation signals in rodent visual cortex. Curr. Opin. Neurobiol. 67, 163–173 (2021).

38. Rodgers, C. C. et al. Sensorimotor strategies and neuronal representations for shape discrimination. Neuron 109, 2308–2325.e10 (2021).

39. Audette, N. J., Zhou, W. X., La Chioma, A. & Schneider, D. M. Precise movement-based predictions in the mouse auditory cortex. Current Biology 32, 4925–4940.e6 (2022).

40. Wekselblatt, J. B., Flister, E. D., Piscopo, D. M. & Niell, C. M. Large-scale imaging of cortical dynamics during sensory perception and behavior. J. Neurophysiol. 115, 2852–2866 (2016).

41. Gerfen, C. R., Paletzki, R. & Heintz, N. GENSAT BAC cre-recombinase driver lines to study the functional organization of cerebral cortical and basal ganglia circuits. Neuron 80, 1368–1383 (2013).

42. Gong, S. et al. Targeting Cre recombinase to specific neuron populations with bacterial artificial chromosome constructs. Journal of Neuroscience vol. 27 9817–9823 Preprint at 10.1523/JNEUROSCI.2707-07.2007 (2007).

43. Madisen, L. et al. Transgenic mice for intersectional targeting of neural sensors and effectors with high spec-ificity and performance. Neuron 85, 942–958 (2015).

44. Bimbard, C. et al. Behavioral origin of sound-evoked activity in mouse visual cortex. Nat. Neurosci. 26, (2023).

45. Harris, K. D. A Shift Test for Independence in Generic Time Series. https://arxiv.org/abs/2012.06862 (2020).

46. Okun, M. et al. Diverse coupling of neurons to populations in sensory cortex. Nature 521, (2015).

47. Stringer, C., Pachitariu, M., Steinmetz, N., Carandini, M. & Harris, K. D. High-dimensional geometry of pop-ulation responses in visual cortex. Nature 571, (2019).

48. Siegle, J. H. et al. Reconciling functional differences in populations of neurons recorded with two-photon imaging and electrophysiology. Elife 10, (2021).

49. Luczak, A. & MacLean, J. N. Default activity patterns at the neocortical microcircuit level. Front. Integr. Neu-rosci. 6, 20722 (2012).

50. Dipoppa, M. et al. Vision and Locomotion Shape the Interactions between Neuron Types in Mouse Visual Cortex. Neuron 98, 602–615.e8 (2018).

51. Burgess, C. P. et al. High-Yield Methods for Accurate Two-Alternative Visual Psychophysics in Head-Fixed Mice. Cell Rep. 20, 2513–2524 (2017).

52. Steinmetz, N. A., Zatka-Haas, P., Carandini, M. & Harris, K. D. Distributed coding of choice, action and en-gagement across the mouse brain. Nature 576, 266–273 (2019).

53. Nestvogel, D. B. & McCormick, D. A. Visual thalamocortical mechanisms of waking state-dependent activity and alpha oscillations. Neuron 110, 120–138.e4 (2022).

54. Syeda, A. et al. Facemap: a framework for modeling neural activity based on orofacial tracking. Nat. Neurosci. 27, 187–195 (2024).

55. Pologruto, T. A., Sabatini, B. L. & Svoboda, K. ScanImage: Flexible software for operating laser scanning mi-croscopes. Biomed. Eng. Online 2, (2003).

56. Pachitariu, M. et al. Suite2p: beyond 10,000 neurons with standard two-photon microscopy. bioRxiv https://doi.org/10.1101/061507 (2016) doi:10.1101/061507.

57. Rupprecht, P. et al. A database and deep learning toolbox for noise-optimized, generalized spike inference from calcium imaging. Nat. Neurosci. 24, (2021).

58. Speed, A., Del Rosario, J., Burgess, C. P. & Haider, B. Cortical State Fluctuations across Layers of V1 during Visual Spatial Perception. Cell Rep. 26, (2019).

